# Online versus Cognitive Control: A Dividing Line between Physical Action and Motor Imagery

**DOI:** 10.1101/2022.10.31.514494

**Authors:** Marie Martel, Scott Glover

## Abstract

Recent work in our lab has shown that motor imagery is highly sensitive to tasks that interfere with executive resources, whereas physical actions are largely immune. This has been taken as support for the Motor-Cognitive model of motor imagery and in opposition to the theory of Functional Equivalence. Here, we examined another prediction of the MCM, namely that an opposite pattern of effects would be observed when the information available for online control was reduced, with physical actions being affected but motor imagery being largely resistant. This was tested in four experiments in which participants performed either physical actions or motor imagery, and in a replication in which they performed both. The experiments manipulated the quality of information available during the online control of movement through: 1) comparing movements made with or without visual feedback (Exp 1 and 1a); 2) comparing movements made using foveal vs. peripheral vision (Exp 2); and 3) comparing physical to mimed actions (Exp 3). All four experiments found evidence in favour of the Motor-Cognitive model in that manipulations of online control affected physical action much more than they affected motor imagery. These results were, however, inconsistent with a Functional Equivalence view. We discuss these results in the broader context of other theoretical views of motor imagery.

**Public Significance Statement:** Motor imagery is a vital component of elite motor skill training and has great utility as a rehabilitative tool in those suffering from brain imagery. A sophisticated understanding of motor imagery is thus critical to optimize its use in these contexts. However, presently there is a paucity of theoretical accounts of motor imagery. A currently popular view is that motor imagery is functionally equivalent to physical actions, yet this account is an oversimplification. We propose instead the Motor-Cognitive model, which incorporates the differences as well as similarities between motor imagery and physical action. Here, we tested the Motor-Cognitive model against the Functional Equivalence view in four experiments that manipulated the amount and/or quality of online visual feedback available and found clear evidence for a difference in motor imagery versus physical action. These experiments provided convincing support that motor imagery is functionally distinct from physical action. This advance in theory regarding motor imagery has important implications for its implementation as both a performance-enhancing and therapeutic tool.

## Introduction

The human ability to generate mental images has fascinated philosophers and psychologists for centuries (Dennett, 1969; Galton, 1880; Hobbes, 1651; Pearson, 2019). Motor imagery represents a specific form of mental imagery in which an action is simulated internally without any associated body movement. In the present study, we tested the Motor-Cognitive model (Glover et al., 2020; Glover & Baran, 2017; Martel & Glover, 2023) against the Functional Equivalence view (Jeannerod, 1995; Jeannerod & Decety, 1995).

In the Motor-Cognitive framework, both physical and imagined movements consist of a planning and execution stage (Glover, 2004). In the planning stage, motor imagery and physical actions share the same internal representations (Jeannerod & Decety, 1995). Information originating from the environment and the body are combined with stored memories of actions to select an appropriate motor plan and form a forward model (for motor imagery: Glover & Dixon, 2013; Kilteni et al., 2018; Vry et al., 2012); and motor control (Glover, 2004; Shadmehr & Krakauer, 2008; Wolpert et al., 1998)). During execution, however, motor imagery and physical action bifurcate.

In a physical action, sensory predictions from the forward model are combined with online feedback; when these diverge, an error signal is sent, and online control is used to correct the movement in flight (Goodale et al., 1986; Paulignan et al., 1991; Pisella et al., 2000; Shadmehr & Krakauer, 2008; Wolpert et al., 1998). Conversely, in motor imagery, the stationary condition of the body results in an absence of feedback-based online control. Rather, monitoring and elaboration of the motor image is held to be a cognitive process that depends heavily on executive functions subserved by the dorsolateral prefrontal cortex (Glover et al., 2020; Glover & Baran, 2017; Martel & Glover, 2023). Unlike the fast visuomotor modules that underlie online control (Goodale et al., 1986; Milner & Goodale, 2006; Paulignan et al., 1991), executive resources are unequipped to simulate the fast, fine-tuned adjustments that occur during physical movement (Nair et al., 2003), and as a consequence, motor imagery becomes less accurate in simulating the characteristics of physical action. This is particularly the case under conditions in which the process of online control is highly engaged during the physical action. For example, when one imagines catching a ball on a windy day, motor imagery should struggle greatly to accurately simulate the large number of online adjustments that would be made during the action. Conversely, when a physical action is ballistic, such as for a fast striking or tapping motion, motor imagery ought to be able to simulate it with a high degree of fidelity.

The Motor-Cognitive model can be contrasted with the most prominent current theory of motor imagery, known as “Functional Equivalence.” This view posits that imagined movements are little more than the internal physiological and neural correlates of physical actions while physical action is inhibited (Jeannerod, 1995; Jeannerod & Decety, 1995). The notion of Functional Equivalence drew early support from findings that motor imagery and physical actions often share timing similarities and biomechanical constraints. For example, movements of greater duration generally also take longer to imagine (Baccarini et al., 2014; Boulton & Mitra, 2013; Decety & Jeannerod, 1995; Decety & Michel, 1989; Frak et al., 2001; Glover et al., 2005; Guillot & Collet, 2005; Jeannerod & Frak, 1999; Macuga & Frey, 2014; Papaxanthis et al., 2002; Roberts et al., 2019). Further, physiological responses such as changes in heart rate are experienced similarly during physical and imagined movement (Decety et al., 1991). Finally, large amounts of neural overlap between motor imagery and physical action have been observed in neuroimaging studies (see for review, Gerardin et al., 2000; Grèzes & Decety, 2001; Hardwick et al., 2018; Hétu et al., 2013; Jeannerod, 2008; Lotze & Halsband, 2006; Macuga & Frey, 2012).

However, numerous behavioural studies have observed large discrepancies between the characteristics of physical actions and motor imagery that are inconsistent with the idea of Functional Equivalence but can be explained within the Motor Cognitive framework (Calmels & Fournier, 2001; Cerritelli et al., 2000; Decety et al., 1989; Gandrey et al., 2013; Glover et al., 2020; Glover & Baran, 2017; Grealy & Shearer, 2008; Hanyu & Itsukushima, 2000; Martel & Glover, 2023; Maruff et al., 1999; Slifkin, 2008; Wong et al., 2013; Yoxon et al., 2015, 2017). Further, at the neural level, certain brain areas show greater activation during motor imagery than during physical action, most notably the dorsolateral prefrontal cortex (Dodakian et al., 2014; Gardini et al., 2016; Gerardin et al., 2000; Guillot et al., 2008, 2009; Lotze et al., 2003; Macuga & Frey, 2014; Stephan et al., 1995; Tacchino et al., 2017; Vry et al., 2012; Zapparoli et al., 2013), and connectivity patterns differ between physical and imagined action (Gao et al., 2011; Kim et al., 2018; Solodkin et al., 2004; Xu et al., 2014).

As stated, an important component of the Motor Cognitive model is the bifurcation of physical action and motor imagery processes during online control. Critically, motor imagery cannot use online feedback to adjust the movement in flight as occurs during physical execution. Instead, motor imagery relies heavily on executive resources to elaborate the motor image. This often results in timing discrepancies between real and imagined actions (Glover et al., 2020; Glover & Baran, 2017; Martel & Glover, 2023; Maruff et al., 1999; Yoxon et al., 2015). For example, performing a demanding executive task, such as counting backwards by threes or generating words from a single letter, greatly slows the execution of motor imagery while only marginally affecting the timing of its physical counterpart (Glover et al., 2020; Glover & Baran, 2017; Martel & Glover, 2023). Further, in line with the proposed role of executive functions in the Motor-Cognitive model, Martel and Glover (2023) observed that motor imagery was slowed by disruption of the dorsolateral prefrontal cortex with TMS, but no such effect of TMS was observed on physical actions.

Another critical prediction of the Motor-Cognitive model alluded to above, is that only physical actions should be sensitive to factors that impact online control, which itself depends on the quality of information available as the movement unfolds. For example, if visual feedback is removed during a movement, physical action will slow to make greater use of proprioceptive feedback (Berthier et al., 1996; Kuhtz-Buschbeck et al., 1999; Martel & Glover, 2023; Sarlegna & Mutha, 2015; Woodworth, 1899). More generally, in circumstances in which the quality of information available to online control is reduced, there will be increased movement times, an increased number of in-flight adjustments, and an extended deceleration phase (Marteniuk et al., 1987; Meyer et al., 1988). However, within the Motor-Cognitive model, motor imagery relies on executive resources rather than online control processes during execution. As a consequence, motor imagery should be impervious to these same effects, at least during execution. This prediction is wholly distinct from one based on the central tenet of the Functional Equivalence view whereby both physical and imagined actions operate using the same internal processes. On that analysis, motor imagery and physical actions should be similarly affected by any and all extraneous variables.

Whereas previous tests of the Motor Cognitive model focussed on comparing the effects of manipulations of executive functions on motor imagery and physical actions (Glover et al., 2020; Glover & Baran, 2017; Martel & Glover, 2023), the present study focussed on the manipulation of online control, and predicts the opposite pattern of effects to those observed previously. Specifically, whereas disrupting execution functions slowed motor imagery but left physical actions largely unaffected, in the present study, we predicted that disrupting online control would slow physical actions but leave motor imagery largely unaffected.

We here manipulated the quality of information available to online control in four experiments. In each experiment, participants were required to either execute or imagine a reaching and placing movement under different conditions. In Experiment 1, participants performed the task either with full vision or no vision during execution. In Experiment 1a the same conditions were repeated within-participants to examine the effects of previous practice with one behaviour on performance with the other. Experiment 2 tested the effects of reaching under foveal vs. peripheral vision, whereas Experiment 3 compared performance when interacting with a real object vs. miming the same movement. In the latter-named conditions of each experiment, online control would be expected to be affected, as the information available to online processes would be reduced. For example, in Experiments 1 and 1a, removing vision coincident with movement onset would eliminate the use of online visual feedback of the hand and target, and thus should impair normal online control processes. In Experiment 2, moving in peripheral vision reduces the visual acuity which again should cause online control to be impaired. In Experiment 3, the lack of physical interaction with the target object in the mime condition should impair online control.

If the Motor-Cognitive model is correct, in each experiment we ought to observe differences between physical actions and motor imagery. Specifically, whereas physical actions ought to be highly sensitive to the variables impacting online control, motor imagery should be comparatively immune. As such, the Motor-Cognitive model predicted an interaction between the Action variable (physical vs. imagined actions) and the respective online control variables in each experiment. Further, any main effects whereby motor imagery took either more or less time than physical actions would be consistent only with the Motor-Cognitive model and not the Functional Equivalence view. Conversely, the only effect allowed under the Functional Equivalence model would be a main effect of online control that should be similar for both motor imagery and physical action.

### Experiment 1

Experiment 1 explored the effects of removing visual feedback during movement on both physical action and motor imagery. As visual feedback plays a key role during online control (Glover, 2004; Meyer et al., 1988), we expected to see large effects of this manipulation on physical actions. In particular, all of movement times, time spent in deceleration, and the number of online corrections should be greater when feedback is removed versus when it is available. If the Motor-Cognitive model is correct, these effects should be much smaller or even absent for motor imagery, leading to an *Action* (physical action vs. motor imagery) by *Vision* (full vision vs. no vision) interaction. Conversely, if the Functional Equivalence view is correct, both physical action and motor imagery should be equally affected by the removal of vision, and no such interaction should arise. Similarly, only the Motor-Cognitive model predicts a main effect of Action, whereby motor imagery should take longer than physical action.

## Methods

### Transparency and Openness

Sample size and inclusion/exclusion criteria for all four experiments were determined *a priori* and were pre-registered prior to data collection alongside study rationales, procedures, data processing and analyses (https://osf.io/z7u9f). The use of likelihood ratios is a statistical approach based on evaluating the strength of evidence rather than on controlling error rates (as opposed to Neymann-Pearson hypothesis testing). Nevertheless, we determined the sample size as that which would be required to establish an 80% or greater chance of obtaining an adjusted likelihood ratio of at least 10:1 for both the predicted main effects and the interaction. We chose 10:1 as our threshold likelihood ratio as we believe this represents strong evidence for one statistical model over another. Predicted effect sizes for Experiment 1 were based on estimates obtained from pilot data on the same task (n=12, https://osf.io/4ne5k/) with effect sizes adjusted for small samples. Calculations were performed separately for all effects to be tested in the experiment: both the kinematic data (all main effects, based on Hedges *g*) and the main effects (again, based on Hedges *g*) and critical interactions (based on *dz*, *f*, the correlations among repeated measures, and attenuation for both main effects) of the keypress data. Where sample size requirements differed among these, the highest value was taken. Calculations for main effects were done in Excel, as no dedicated software is available to calculate “power” for likelihood ratios. Calculations for interactions were estimated using *dz* for main effects and *f* for the interaction, adjusting for the correlation among repeated measures and the attenuation observed in both the *Action* and *Vision* conditions, and then calculating the sample size that would be needed to obtain a p-value of 0.0135 (roughly corresponding to an adjusted likelihood ratio of 10:1) using Gpower (v. 3.1.9.6). A file showing these calculations is available here: https://osf.io/4ne5k/. All data, code and material used in this study are available on the website of the Open Science Framework (OSF) at the following link: https://osf.io/a9tzf/. Data collection took place from September to October 2021.

### Participants

We tested 32 participants (25 women, 7 men, age range 18-35 years) from among Royal Holloway students and staff, with 16 participants pseudo-randomly assigned to each of the Physical Action and Motor Imagery groups. Two participants were replaced because they failed to follow the instructions. Participants from the undergraduate research participation pool received research credits, others received monetary rewards or no compensation. All participants were right-handed by self-report, had normal or corrected-to-normal vision, had no motor or neurological impairments, and were naïve as to the exact purpose of the experiment. Due to Covid-19 safety precautions, all participants performed all tasks with a face mask and surgical gloves on. This and all other experiments presented here were approved by the College Ethics Board at Royal Holloway.

### Experimental setup

The general experimental setup is shown in Figure 1 and has been used previously (Glover et al., 2020; Glover & Baran, 2017; Martel & Glover, 2023). Participants sat comfortably at a 50 × 90 cm table set inside a large room. A computer keyboard, a wooden box (dimensions: 15.7 × 42 × 22 cm) and a small white plastic “poker chip” disc sat on top of the table. The box contained a grey plastic tube (dimensions: 15 × 4.2 cm) positioned 15 cm from the side of the box close to the participant, and the Polhemus receiver module, positioned in the corner of the box furthest from the participant. The starting position was a yellow square sticker (0.5 × 0.5 cm), positioned 52 cm away from the left side of the table, and 5 cm away from the near side. The disc was 1 cm thick and 4 cm in diameter. All other dimensions and positions were as shown in Figure 1.

**Figure 1.**
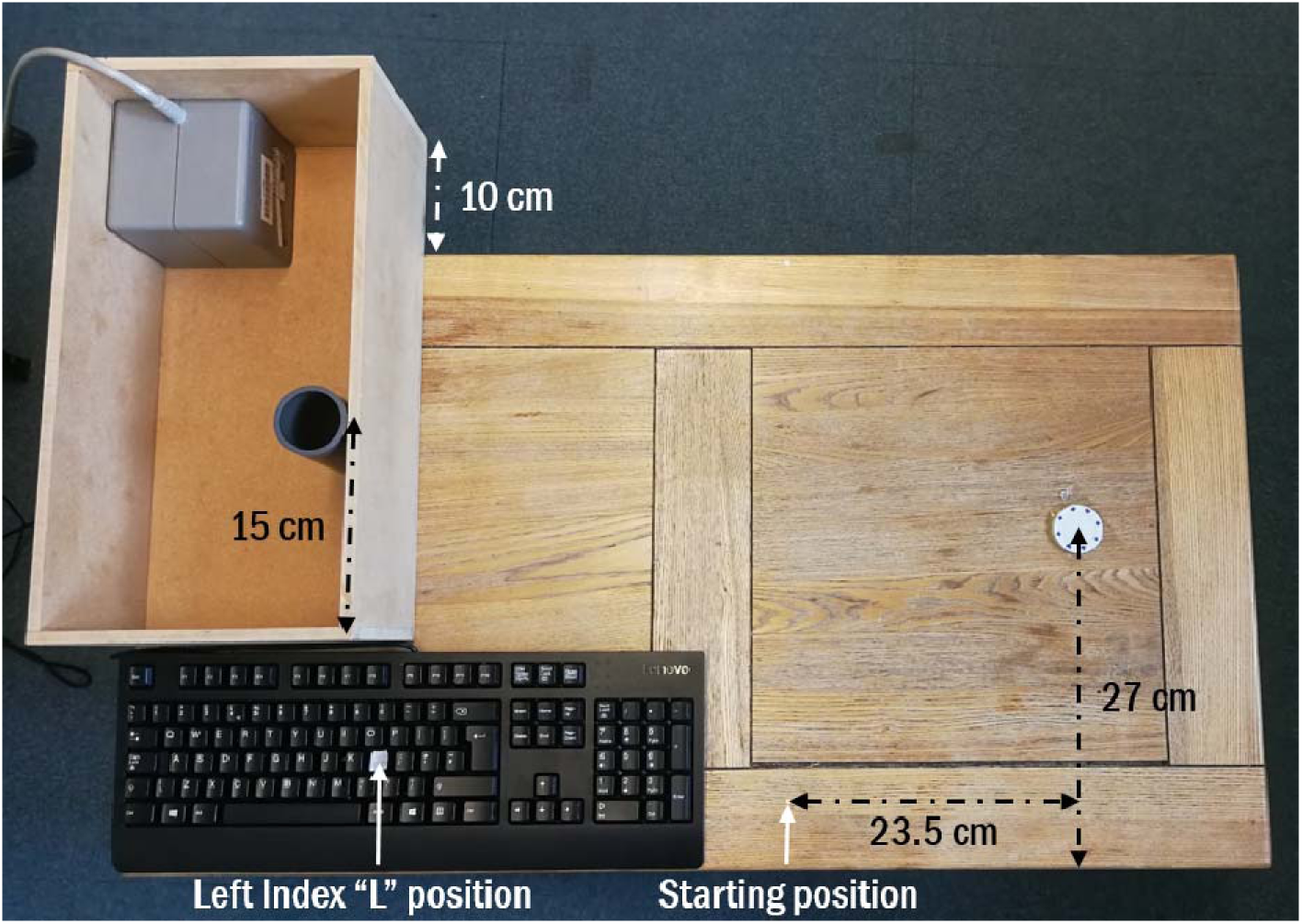
Experimental setup. Participants began with their right thumb and index finger on the starting position, and at a signal had to either reach to grasp the disc then place it into the grey cylinder on the left, or imagine doing so while remaining stationary. Participants used their left index finger to indicate the start and end of the movement (or its imagined counterpart) by pressing the “L” key.

For the participants in the Physical Action group, we placed two Polhemus transmitters (7 mm in diameter) on the participant’s right thumb and index fingernails to record their spatial location in x,y,z coordinates (Polhemus Fastrak system; sampling rate: 120 Hz). Data was analysed offline.

### Procedure

Each trial began with participants sitting with their body midline aligned with the starting mark, their right thumb and index finger in a comfortable “pinch grip” posture on this mark, and their left index finger resting on the “L” key. After a starting tone triggered via PC, participants had to reach for the disc and place it into the cylinder, or imagine doing so. For both the Physical Action and Motor Imagery groups, participants indexed the beginning and the end of their movement by pressing the “L” key with their left index finger. The Physical Action group performed the grasping and placing movement with their eyes either remaining open throughout (Full Vision trials) or after closing their eyes at the onset of each movement (No Vision trials). The experimenter monitored participants to ensure they followed instructions. The Motor Imagery group imagined reaching and placing the disc in the same two visual conditions. Participants had to imagine the action as vividly as possible, using both visual and kinaesthetic imagery of their movement experienced in the first person (i.e., to imagine seeing and feeling themselves performing the movement), while keeping their right arm/hand still on the starting position.

Each participant performed two counterbalanced blocks of 6 practice and 12 experimental trials each for a total of 24 analysed trials (12 Full Vision and 12 No Vision). Trials which had to be excluded from analysis due to participant error were not repeated, resulting in some participants having fewer than 24 accurate trials before data processing. Overall, 26/768 (3.4%) trials were not analysed: 4/192 and 7/192 in the Motor Imagery group, No Vision and Full Vision respectively; 11/192 and 4/192 in the Physical Action group, No Vision and Full Vision respectively. For the Physical Action group trials, the experimenter put the disc back in place and participants returned to the starting position. Once the participant was set, the experimenter triggered the tone to begin the next trial.

### Data processing

#### Keypress data

We extracted reaction time (RT) and movement time (MT) from the keypress data for both Physical Action and Motor Imagery. RT was the time between the starting tone and the first keypress (indicating the beginning of the movement/imagery). MT was measured as the time between the first and second keypress. In addition to the trials removed before analysis, any trial with RT or MT outside ± 2 IQR (Interquartile Range) for each combination of block and participant was removed from the analysis (3.4% of the total trials; Motor Imagery Full Vision: 0.40%; Motor Imagery No Vision: 0.40%; Physical Action Full Vision: 1.1%; Physical Action No Vision: 1.5%). As we did not make any prediction on RT, we only present the data on MT in this report (RT data and analysis for each experiment are presented in the Supplementary Information).

#### Kinematic data

We extracted kinematic indices of each movement using a custom program on Python. We calculated velocity from x,y,z coordinates and filtered the data at 15 Hz with a low pass Butterworth filter of order 2. Each movement was divided into a reaching and placing phase. The beginning of the first, reaching phase was set as the point at which a minimum speed of 5cm/s was obtained, with the reaching component marked as ended at the time when the hand had again gone above a velocity of 5 cm/s after having fallen below it (or reached its minimum value), following the beginning of the first movement. This latter point also defined the beginning of the second, placing phase. The end of the placing phase was defined in the same way as the end of the reaching phase. For each of the placing and reaching phases, we also measured the time spent in deceleration (i.e., the time between the peak velocity and end of each movement phase), and the total number of online adjustments (indexed as the number of re-accelerations of the hand during the deceleration subphases and/or decelerations of the hand during acceleration subphases of each movement). Any trial with RT or MT outside ± 2 IQR (Interquartile Range) for each combination of block and participant was removed from the analysis (3.2% of the total trials; Full Vision: 0.27%; No Vision: 2.97%).

Additionally, data obtained via the Polhemus was used to perform a manipulation check that participants were pressing the “L” key at times closely corresponding to their actual movement onset and offset. We compared RT and MT indicated by the keypresses with kinematically determined RT and three different measures of MT (see Supplementary Information).

### Analyses and statistics

Data were analysed using R (R studio version: 1.4.1717, R Core Team, 2018) and nested linear mixed models (package *lme4,* Bates et al., 2015) with the evidence for competing statistical models calculated using adjusted likelihood ratios (λ_adj_) (Glover & Dixon, 2004). Likelihood ratios represent the relative likelihood of the data given two competing models, providing a simple, intuitive index of the strength of the evidence while avoiding many of the pitfalls associated with NHST (Wasserstein & Lazar, 2016). As models with more parameters will almost always fit the data better than models with fewer parameters, adjusted likelihood ratios (λ_adj_) were computed using the Akaike Information Criterion (Akaike, 1973). Likelihood ratios are closely related to *p*-values, and under typical hypothesis testing conditions, a *p*-value of 0.05 would correspond to a λ_adj_ ∼ 2.7. The use of nested linear models in combination with likelihood ratios allows for direct comparisons between models including multiple independent factors, something not directly possible using *p-*values.

To test the efficacy of our *Vision* manipulation (Full Vision/No Vision), we compared its effects on the kinematic indexes (movement time, total deceleration time, and number of online adjustments). We used a 2×2 mixed design, with *Action* (Physical Action/ Motor Imagery) as a between-participants variable, and *Vision* (Full Vision/No vision) as a within-participants variable, on the movement times as indexed by the keypresses. We implemented the maximum random effects structure for which the respective model still converged and improved model fit (Barr et al., 2013), which included a random slope for each participant when possible. We present analyses of the movement times based on keypresses. Exploratory analyses on reaction times are presented in the Supplementary Information.

We report the *f²* effect size for the keypress, and the *dz* for kinematics, based on t-tests and/or the ANOVA of the full model for each experiment.

## Results

### Kinematic indices

Table 1 shows the effects of *Vision* on the kinematic indices in the Physical Action group reaching and placing task. As expected, performing the movement without vision greatly increased movement times, time spent in deceleration, and the number of online adjustments for both the reaching and placing phases of physical actions (all λ_adj_> 1000, *dz* range between 2.4 and 6.7). We concluded from this that our manipulation of online control had the intended effects on physical actions. Additionally, there was very little difference between movement times when comparing the analyses of kinematics and the keypress data, suggesting that participants were overall highly accurate at indicating the beginning and end of their movements. This was shown in an additional analysis performed on kinematic MT for the physical action group vs. keypress MT for the motor imagery group which rendered the same results as in the next section (see Supplementary Information), confirming the results were not simply an artefact of the use of keypresses to index movement times in the physical action group.

**Table 1.** Effect of moving with full vision or no vision on kinematics.

| | Full Vision (mean ± SD) | No Vision (mean ± SD) |
| --- | --- | --- |
| Total movement time (ms) | 1980 ± 365 | 4400 ± 1145 |
| **Reaching phase** |  |  |
| Deceleration Time (ms) | 637 ± 153 | 1331 ± 366 |
| Number of online adjustments | 1 ± 0.7 | 4.5 ± 1.6 |
| **Placing phase** |  |  |
| Deceleration Time (ms) | 743 ± 156 | 2437 ± 736 |
| Number of online adjustments | 1.7 ± 0.8 | 9.2 ± 2.9 |

### Keypress movement times

Figure 2 shows the mean keypress movement times as a function of *Vision (Full Vision/No Vision)* and *Action (Physical Action/Motor Imagery groups)* conditions. The best-fitting model of the keypress MT data included the effect of *Vision* and the *Vision* × *Action* interaction. A statistical model including an effect of *Action* only did not fit the data noticeably better than the null model (λ_adj_ = 1.8, f² = .008). However, adding the effect of *Vision* improved the fit greatly (λ_adj_ > 1000, f² = 1.02), with longer movement times in the Physical Action group, and adding the interaction *Action* × *Vision* improved the fit even further (λ_adj_ > 1000, f² = .94), reflecting increased movement times in the No Vision condition but only for the Physical Action group. To confirm this, we ran an additional analysis only on the MI group: here the evidence modestly favoured the null model over a model that included an effect of *Vision* (λ_adj_ = 2.63, *dz* = 0.035), suggesting that for the Motor Imagery group, MT remained consistent whether performed in the Vision or No Vision condition. Additionally, to further explore whether the lack of *Action* effect was, as we contended, driven by the increased MT in the No Vision condition for the Physical Action Group, we investigated the presence of the *Action* effect in the Full Vision condition only. The evidence showed greater MT for the Motor Imagery Group compared to the Physical Action Group (λ_adj_ = 140, *dz* = 0.67).

**Figure 2.**
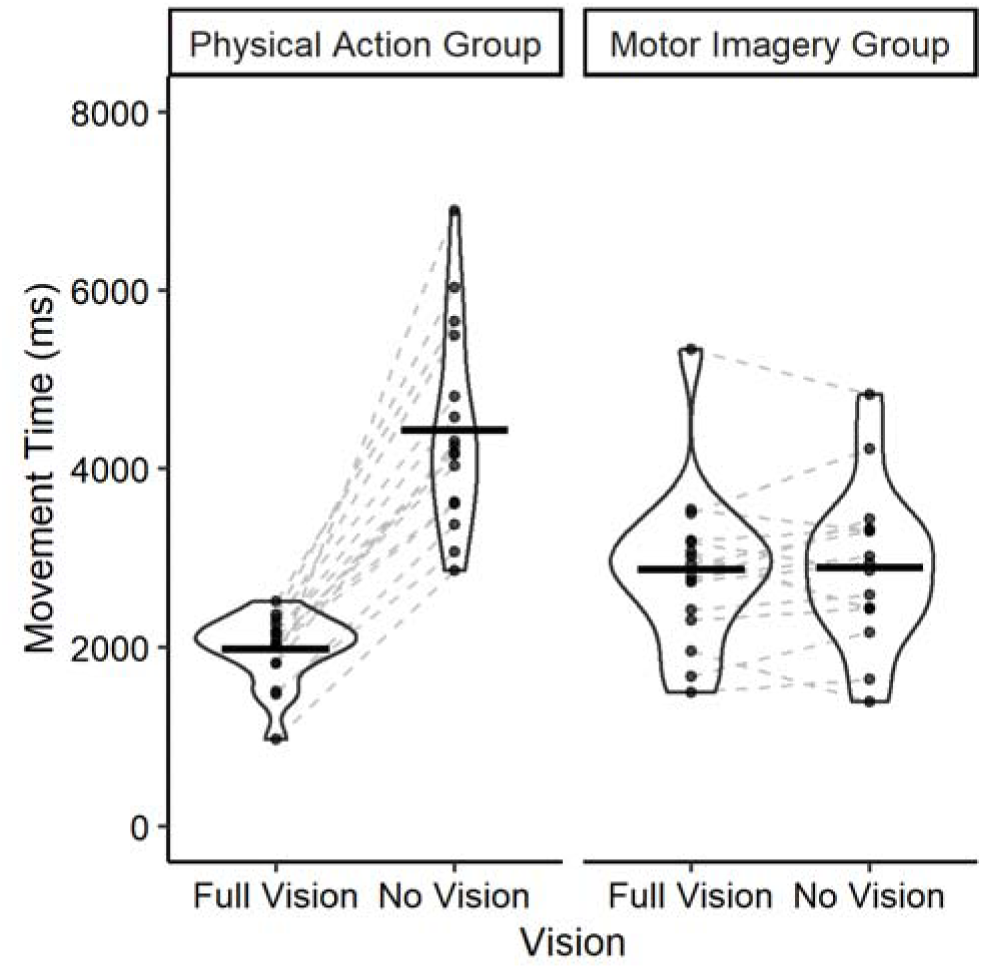
Effects of varying vision (Full Vision vs. No Vision) on movement times in the Physical Action and Motor Imagery groups. Individual points represent the mean score for single participants, dashed grey lines pair the means of individual participants together, and horizontal lines represent the group means. Enclosures represent the distribution of the presented data, with wider horizontal sections indicating a larger number of observations.

To directly assess the Motor-Cognitive and Functional Equivalence views, we compared statistical models incorporating the predictions made by each. A full model including both main effects and their interaction, as predicted by the Motor-Cognitive model, fit the data much better than a model including only an effect of *Vision*, as predicted by the Functional Equivalence model (λ_adj_ > 1000). In other words, the data were over 1000 times as likely when including the effects uniquely predicted by the Motor-Cognitive model than when assuming only the effect predicted by the Functional Equivalence view.

## Discussion

In Experiment 1, the kinematic analysis confirmed that our manipulation was successful. All three indices of online control in physical actions, including movement times, the number of online adjustments, and the time spent in deceleration, were much greater when vision was removed than when it was available. The effects were most pronounced in the placing phase of the movement, which required more precision than the grasping phase. This showed that the movement times for physical actions were related to the amount of information available to online control.

The keypress data showed that whereas there were large effects of removing vision on physical actions, imagined movement times were not different in vision and no vision conditions. These contrasting effects of vision on imagery and physical action were consistent with the Motor Cognitive view, whereby physical actions should be more affected than motor imagery by manipulations that impact online control. They were not, however, compatible with a Functional Equivalence view, wherein both physical actions and motor imagery share common internal processes and so should be equally affected by extraneous variables. Interestingly, though we did not conduct post-testing interviews as a matter of course, many participants in the motor imagery group reported that they had imagined performing the movements quite straightforwardly in both the vision and no vision conditions, consistent with the near-identical times for the two vision conditions in the keypress data of the motor imagery group. This was in stark contrast to the numerous difficulties those in the physical action group invariably experienced in the no vision condition relative to the vision condition.

Surprisingly at first glance, the predicted main effect of *Action* was not present. Although physical actions in the vision condition took significantly less time than their imagined counterparts – itself a typical finding in this task (Glover et al., 2020; Glover & Baran, 2017; Martel & Glover, 2023), the same movements in the no vision condition took less time in the motor imagery group than in the physical action group. In short, it would appear that the *Vision × Action* interaction obfuscated a potential main effect of *Action*. Our interpretation of these results was that the effect of removing vision on physical action was so large that movement times across both conditions averaged out to be approximately the same, and thus a main effect of *Action* was not observed.

### Experiment 1a

Experiment 1a provided a replication of Experiment 1, but added an *Order* condition using a within-participants design to measure the effects of having physically practiced the task on motor imagery performance, and vice-versa. In past research, it has been observed that physical practice can lead to a closer correspondence between physical action and motor imagery movement times (Wong et al., 2013; Yoxon et al., 2015, 2017). One possibility therefore is that lack of experience with the Physical Action version of the task in the Motor Imagery group, and in particular in the no vision condition, was responsible for the large interaction between *Vision* and *Action*. This may have resulted because participants in the Motor Imagery group might not have been able to accurately imagine the movement in the absence of vision, due to not having experienced it before. Although effects of physical experience of the task on motor imagery would be consistent with the Motor Cognitive model, as motor imagery is held to depend to a significant extent on the strength of the internal representations of actions (which can be strengthened by physical experience), and we have reported such effects before (Martel & Glover, 2023), we nonetheless sought to examine whether such practice effects might be sufficient to eliminate the interaction between the *Action* and *Vision* variables by making motor imagery movement times more similar to physical action.

Here, participants performed both the Physical Action and Motor Imagery conditions of the same task as in Experiment 1, with half taking part in the physical action conditions first, followed by the motor imagery task, and the other half performing the tasks in the opposite order. Both groups performed each task under *Vision* and *No Visio*n conditions.Whereas it might be expected that performing the Physical Action task first would reduce the difference in movement times between Physical Action and Motor Imagery, the Motor Cognitive model nevertheless predicts the same general pattern of results as in Experiment 1: physical actions should be more affected than motor imagery by the removal of visual feedback in the No Vision condition. In contrast, the Functional Equivalence view holds that there should again be no overall differences in movement times between Physical Action and Motor Imagery tasks, with both equally affected by the removal of vision.

## Methods

### Participants

We tested 32 participants (24 women, 8 men, tested age range 17-27 years) from Royal Holloway students. Three participants were replaced because of technical errors during recording. Inclusion criteria were the same as in Experiment 1. During this experiment, there were no Covid-19 restrictions so participants did not wear masks or gloves. Sample size and inclusion/exclusion criteria were determined *a priori* as per Experiment 1 and were pre-registered prior to data collection alongside study procedures and data processing and analyses (https://osf.io/2dmqz). The full dataset and scripts are available to download at the following link: https://osf.io/a9tzf/. Data collection took place from June to December 2024.

### Experimental setup and procedure

Setup, starting position, instructions and number of trials were the same as in Experiment 1. Out of 1536 trials in total, 138 (9.9%) were excluded and not repeated due to participant errors (Motor Imagery Full Vision: 13/384; Motor Imagery No Vision: 14/384; Physical Action Full Vision: 32/384; Physical Action No Vision: 79/384). The experiment was identical to Experiment 1 except that all participants took part in both Physical Action and Motor Imagery conditions, with the order of conditions counterbalanced.

### Data processing

Data processing was performed exactly as in Experiment 1. In the keypress data, 4.8% of the trials had RT or MT outside ± 2 IQR (Interquartile Range) for each combination of block and were removed (Motor Imagery Full Vision: 1.1%; Motor Imagery No Vision: 1.0%; Physical Action Full Vision: 1.5%; Physical Action No Vision: 1.2%). In the kinematic data, the number of trials removed was 5.9% (Full Vision: 3.2%; No Vision: 2.7%). Correlations between keypress and kinematic data, as well as exploratory analyses on RT can be found in the Supplementary Information.

### Analyses and statistics

We used a 2×2×2 mixed design, with *Order* (Physical Action First/Motor Imagery First) as a between-subjects variable, and *Vision* (Full Vision/No vision) and *Action* (Motor imagery/Physical action) as within-subjects variables, on the movement times as indexed by the keypresses. Everything else was as in Experiment 1.

## Results

### Kinematic indices

Table 2 shows the effects of *Vision* on the kinematic indices in the physical reaching and placing task. As expected, performing the movement in the no vision condition resulted in much greater movement times, time spent in deceleration and online adjustments of both the reaching and placing phases of physical actions (all λ_adj_> 1000, *dz* range between 1.4 and 5.0). We concluded from this that our manipulation of online control again had the intended effects on physical actions. Additionally, there was very little difference between movement times when comparing the analyses of kinematics and the keypress data, suggesting that participants were overall highly accurate at indicating the beginning and end of their movements. This was shown in an additional analysis performed on kinematic MT for the physical action condition vs. keypress MT for the motor imagery condition, which rendered similar results as in the next section (see Supplementary Information), confirming the results were not simply an artefact of the use of keypresses to index movement times in the physical action condition.

**Table 2.**
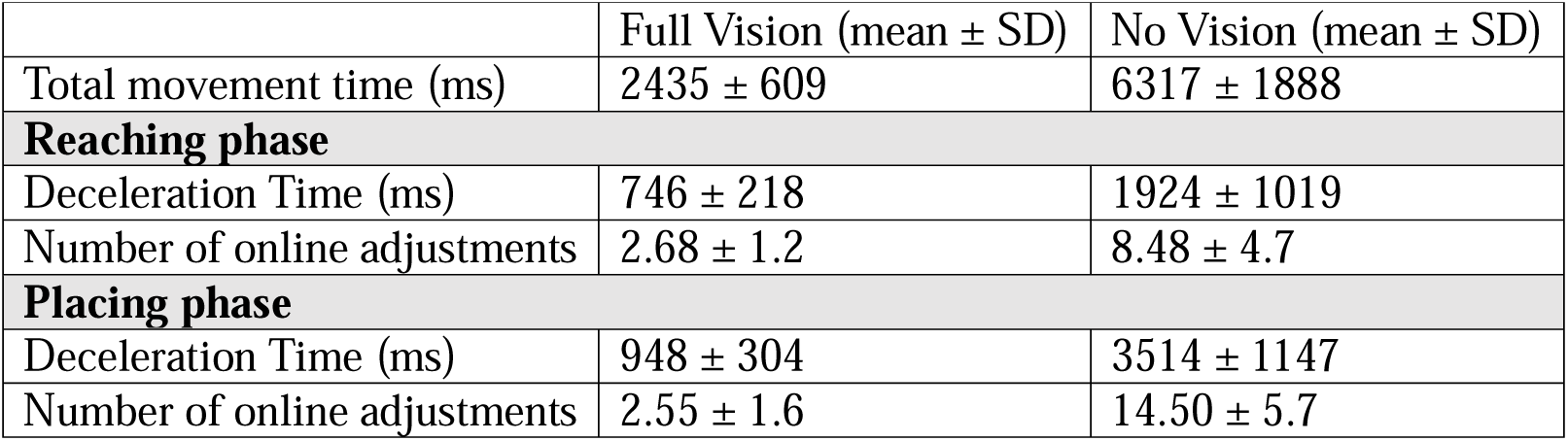
Effect of moving with full vision or no vision on kinematics.

| | Full Vision (mean ± SD) | No Vision (mean ± SD) |
| --- | --- | --- |
| Total movement time (ms) | 2435 ± 609 | 6317 ± 1888 |
| **Reaching phase** |  |  |
| Deceleration Time (ms) | 746 ± 218 | 1924 ± 1019 |
| Number of online adjustments | 2.68 ± 1.2 | 8.48 ± 4.7 |
| **Placing phase** |  |  |
| Deceleration Time (ms) | 948 ± 304 | 3514 ± 1147 |
| Number of online adjustments | 2.55 ± 1.6 | 14.50 ± 5.7 |

**Table 3.** Effect of using foveal or peripheral vision on kinematics.

| | Foveal Vision (mean ± SD) | Peripheral Vision (mean ± SD) |
| --- | --- | --- |
| Total movement time (ms) | 1863 ± 344 | 2943 ± 782 |
| **Reaching phase** |  |  |
| Deceleration Time (ms) | 606 ± 131 | 842 ± 205 |
| Number of online adjustments | 0.9 ± 0.6 | 2.4 ± 1.1 |
| **Placing phase** |  |  |
| Deceleration Time (ms) | 732 ± 182 | 1576 ± 524 |
| Number of online adjustments | 1.3 ± 1.1 | 5.4 ± 2.5 |

### Keypress movement times

Figure 3 shows the mean keypress movement times as a function of *Vision*, *Action* and *Order* conditions. The best-fitting model of the keypress MT data included a main effect of *Action*, *Vision* and their interaction. A statistical model including only an effect of *Action* whereby there were longer movement times in the physical action condition fit the data much better than a null model (λ*_adj_* > 1000, *f²* = .16). Adding *Vision* to the model vastly improved the fit (λ*_adj_* > 1000, *f²* = .41), with longer movement times in the No Vision condition, and adding the interaction *Action* × *Vision* improved the fit even further (λ_adj_ > 1000, *f²* = 1.3), suggesting that MTs were more affected by Vision in the Physical Action than the Motor Imagery condition.

**Figure 3.**
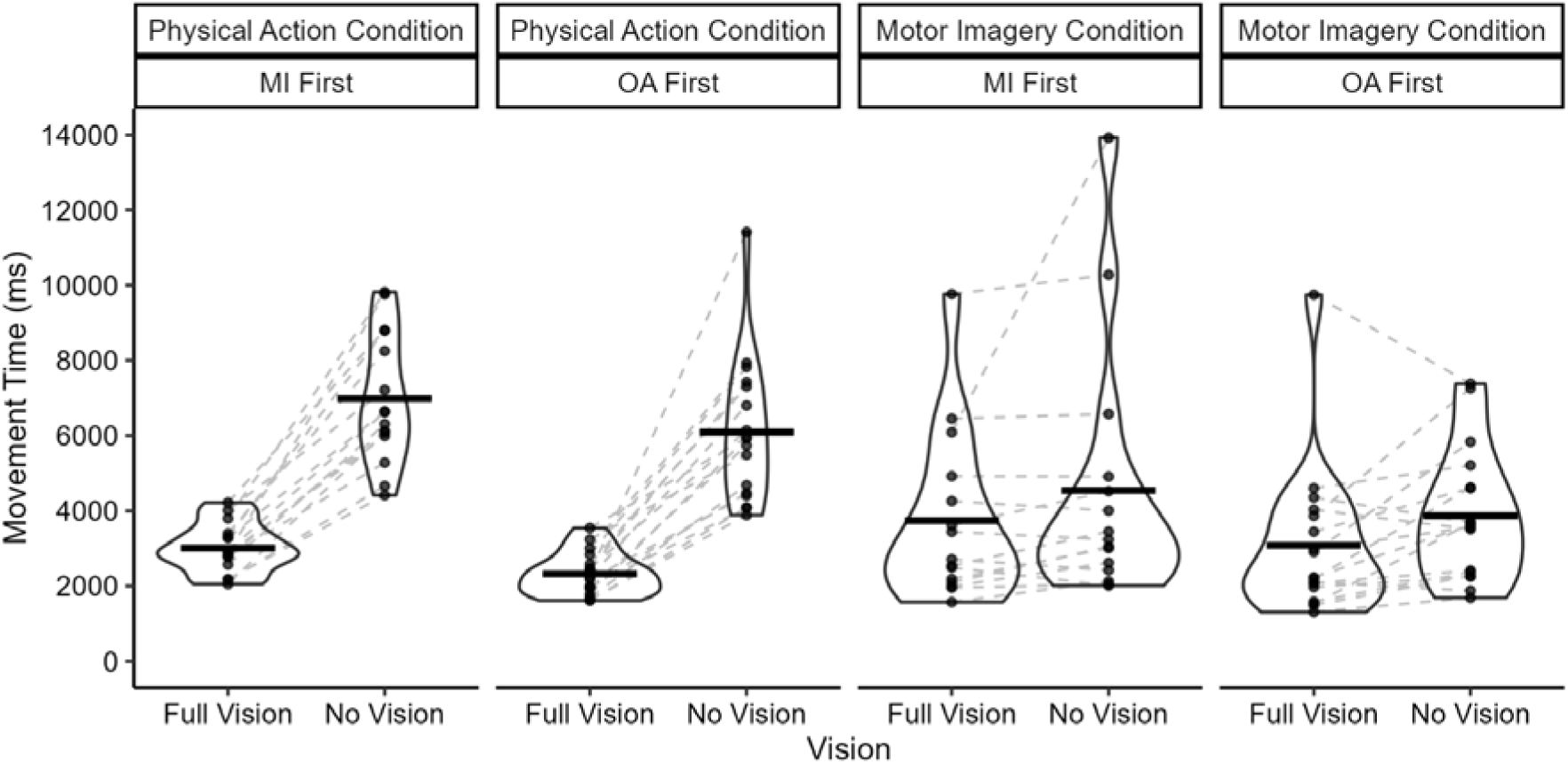
Effects of varying vision (Full Vision vs. No Vision) on movement times in the Physical action and Motor imagery conditions, according to the Order of the blocks. Other conventions as in Figure 2.

The fit was not further improved by adding any of the effects of *Order* or their interactions; rather the evidence suggested that adding these variables made the fit worse (λ_adj_ = 3.4 in favour of the model excluding the effect of *Order* and its interactions, f*²* = .01 for the model including these effects). As in Experiment 1, we conducted an additional analysis of the effects of Vision in the MI condition alone. Here, the best-fitting model included an effect of Vision (λ*_adj_* = 10.2, *f²* = 4.21), suggesting that Motor Imagery MT was affected by the manipulation but to a much lesser extent than Physical Action (mean difference between MT for NV vs. V: 797 ms for Motor Imagery and 3862 ms for Physical Action). The fit was not improved by the addition of *Order* or the interaction between *Order* and *Vision (*λ*_adj_* = 4.95 in favour of the model excluding these effects*, f²* = .09 for the model including these effects).

Finally, to directly assess the Motor-Cognitive and Functional Equivalence views, we compared statistical models incorporating the predictions made by each. A model including the main effects of *Action* and *Vision* and their interaction, as predicted by the Motor Cognitive model, fit the data much better than a model including only an effect of *Vision*, as predicted by the Functional Equivalence model (λ_adj_ > 1000).

## Discussion

Experiment 1a replicated the main results from Experiment 1 and further supported the Motor Cognitive model over the Functional Equivalence view. Kinematic data showed that the manipulation was again successful in affecting physical actions, with longer movement times, longer deceleration times, and more online corrections in the No Vision condition than in the Vision condition. The keypress data showed that whereas there were very large effects of removing vision on physical actions, an interaction existed whereby its effects on motor imagery were much smaller.

Unlike Experiment 1, we here did observe an effect of *Action*, with movement times being greater overall in the Physical Action condition than in the Motor Imagery condition. This was a result of the greater effects of removing vision on the former than on the latter, as confirmed by the additional analysis on imagined and physical movement times under full vision. Although we did not predict such an effect owing to it not being observed in Experiment 1, it was not entirely surprising and the overall pattern of results was again consistent with the Motor Cognitive model but not the Functional Equivalence view.

Of interest was the complete absence of any evidence for the effects of *Order* or its interactions with other variables. This finding was unexpected, as we anticipated observing at least some attenuation of the difference between motor imagery and physical action in the No Vision condition for participants who first experienced the physical action task. This notion was based on past research showing physical practice led to a narrowing of the differences between motor imagery and physical action (Martel & Glover, 2023; Wong et al., 2013; Yoxon et al., 2015, 2017). One possible explanation for this discrepancy is that the small number of trials in the present study was not sufficient practice for participants to incorporate their experiences of the physical action into their planning of motor imagery. This explanation seems unlikely, however, given we did observe a reduction in the difference between Motor Imagery and Physical Action in a previous study with a slightly smaller number of trials (15 per condition in Martel and Glover, 2023; vs. 18 here).

A second explanation for the lack of *Order* effects, and one we favour, is that because the effects of removing vision are concentrated in the online control phase of the movement, motor imagery was unable to incorporate them. As such, despite direct physical experience showing that movements take much longer and generally require a much greater number of online corrections when vision is removed, participants nonetheless imagine these same movements in essentially the same way as they do when no physical practice is given: the motor image is constructed as if the movement were being performed straightforwardly and without the need for online adjustments. This latter hypothesis was consistent with verbal reports offered by many participants: All who commented reported that they executed the No Vision task in the motor imagery condition as if they were directly reaching to and grasping the disc and directly placing it into the cylinder, without any of the fumbling that often occurred in the same condition during the physical action version of the task.

It is also notable that, unlike in Experiment 1, there was evidence for a Vision effect in the motor imagery condition alone (though the effect was nearly five times greater in the physical action condition). At first glance, this seems to contradict the Motor Cognitive model. However, we note that within the model the motor plan that forms the basis of both motor imagery and physical action would likely incorporate a need to slow down the movement in the No Vision condition. This would lead to a movement of lower overall velocity and concomitant greater movement times in both conditions. However, as the majority of the effect of removing vision arises during the online control phase the greatest effects of vision should be on physical action. All that said, the same effect of Vision on motor imagery was not evident in Experiment 1, and thus said effect appears to be unreliable.

### Experiment 2

Experiment 2 provided a further test of the effects of manipulating online control on motor imagery and physical actions. Here, participants performed (or imagined performing) the movement while either being allowed to freely move their eyes and thus foveate the target (foveal condition), or while maintaining fixation on the starting point and conducting the entire movement after leaving the starting position in the visual periphery (peripheral condition). Because of the greater receptor density in the fovea versus the periphery (Jonas et al., 1992), this manipulation ought to have predictable effects on physical action similar to, if not as powerful as, removing vision entirely. Specifically, we expected to see longer movement times, more time spent in deceleration, and a greater number of online adjustments in the peripheral than in the foveal condition. If the Motor Cognitive model is correct, these extended movement times should only arise in the physical action group and not the motor imagery group. Conversely, if the Functional Equivalence view is correct, both physical action and motor imagery should be equally affected by the manipulation.

## Methods

### Participants

We enrolled 60 participants (48 women, 12 men, tested age range 18-44 years old) from Royal Holloway students and staff, 30 participants in each group. Two participants were replaced because of a technical error during recording. Two additional participants were replaced because they failed to follow the instructions. Inclusion criteria and Covid-19 safety precautions were the same as in Experiment 1. Sample size and inclusion/exclusion criteria were determined *a priori* as per Experiment 1 and were pre-registered prior to data collection alongside study procedures and data processing and analyses (https://osf.io/w92aj). The full dataset and scripts are available to download at the following link: https://osf.io/a9tzf/. Data collection took place from November 2021 to March 2022.

### Experimental setup and procedure

Setup, starting position and number of trials were the same as in Experiment 1. Out of 1440 trials in total, 81 (5.6%) were excluded and not repeated due to participant error (Motor Imagery Foveal: 7/360; Motor Imagery Peripheral: 9/360; Physical Action Foveal: 13/360; Physical Action Peripheral: 52/360). Participants in the Physical Action group performed the movement while either being allowed to move their eyes freely to foveate the target and track their hand (Foveal trials), or while being required to maintain fixation on the starting position (Peripheral trials). The experimenter continuously monitored the participants during each trial to ensure they followed instructions. The Motor Imagery group imagined reaching and placing the disc in the same two visual conditions. Participants had to imagine this action as vividly as possible, using both visual and kinaesthetic imagery of their movement experienced in the first person (i.e., to imagine seeing and feeling themselves performing the movement), while keeping their right arm/hand still.

### Data processing

Data processing was performed exactly as in Experiment 1. In the keypress data, 3.2% of the trials had RT or MT outside ± 2 IQR (Interquartile Range) for each combination of block and were removed (Motor Imagery Foveal: 0.66%; Motor Imagery Peripheral: 0.59%; Physical Action Foveal: 0.66%; Physical Action Peripheral: 1.32%). In the kinematic data, the number of trials removed was 5% (Foveal: 1.8%; Peripheral: 3.2%). Correlations between keypress and kinematic data, as well as exploratory analyses on RT, can be found in the Supplementary Information.

### Analyses and statistics

Analyses were the same as in Experiment 1, except that the *Vision* condition was now broken into Foveal/Peripheral conditions rather than Vision/No Vision.

## Results

### Kinematic indices

Table 2 shows the effects of *Vision* on the kinematic indices in the physical reaching and placing task. As expected, performing the movement using peripheral vision increased movement times as well as time spent in deceleration and number of online adjustments in both the reaching and placing phases, compared to foveal vision (all λ_adj_ > 1000, *dz* range between 1.4 and 5.1). Additionally, there was again very little difference between movement times as measured by the kinematics and the corresponding keypresses for each trial, suggesting that participants were highly accurate overall in indicating the beginning and end of the movement when pressing the keys. This was supported by an additional analysis performed on kinematic MT for the physical action group vs. keypress MT for the motor imagery group which rendered the same results as in the next section (see Supplementary Information), confirming the results were not an artefact of the use of keypresses to index movement times in the physical action group.

### Keypress movement times

Figure 3 shows the keypress movement times as a function of *Action* and *Vision* conditions. The best-fitting model of keypress MT was a full model including the effects of *Action*, *Vision* and their interaction. A statistical model including only an effect of *Action* fit the data much better than a null model (λ_adj_ = 803, f² = .042), with longer movement times in the motor imagery group than in the physical action group, consistent with what is typically observed (Glover et al., 2020; Glover & Baran, 2017; Martel & Glover, 2023). Adding *Vision* to the model vastly improved the fit (λ_adj_ > 1000, f² = 3.92), with longer movement times in the Peripheral condition than in the Foveal condition. Finally, adding the interaction *Action* × *Vision* also greatly improved the fit (λ_adj_ = 716, f² = .18): the effect of *Vision* was greater in the physical action group than in the motor imagery group. Similar to Experiment 1a, an effect of *Vision* was present when analysing the Motor Imagery group alone (λ_adj_ = 36, *dz* = 0.58). Thus, whereas motor imagery was not immune to the *Vision* variable in this experiment, the effects of *Vision* were still more than twice as large on physical actions than on motor imagery (1297 ms and 505 ms for the physical action group and motor imagery, respectively). As before, we also directly tested a full model including both the main effects and the interaction, based on the predictions of the Motor-Cognitive theory, to a model including only an effect of *Vision* based on the Functional Equivalence view. The statistical model representing the former was over 1000 times as likely to produce the data than the model predicted by the Functional Equivalence view (λ_adj_ >1000), again offering strong support for the Motor-Cognitive model over the Functional Equivalence view.

## Discussion

It was once again clear from the kinematic data that our manipulation had the desired effect on physical actions: all of the variables including movement time, deceleration time, and number of online adjustments were greater in the peripheral vision condition than in the foveal condition. Thus, we can conclude that the overall increase in movement times in the peripheral condition was due to difficulties in using online control.

In contrast to these effects on physical actions, and consistent with the predictions of the Motor Cognitive model, we observed an interaction between *Action* and *Vision*. However, as in Experiment 1a, we observed a main effect of *Action* whereby movement times were overall greater in the motor imagery than physical action group. These results strongly favour the Motor Cognitive model over the Functional Equivalence view.

Notably, motor imagery was also affected, albeit to a much lesser extent than physical actions, by the requirement to perform the imagined movement while maintaining fixation on the starting position. Although this might again reflect a common planning mechanism for reducing movement velocity affecting both physical actions and motor imagery, another possible explanation in this experiment is that the natural inclination of many participants when performing motor imagery was to follow the path of their imagined movement with their eyes. This was often observed being done by participants in the foveal condition. When prohibited from doing so by the requirement to fixate the starting position, it may be that some participants found it more difficult to imagine the movement and thus slowed down as a result. Unfortunately, we did not record which participants did or did not move their eyes to follow their imagined trajectory during the foveal trials, and so were unable to correlate this behaviour with performance. As such, this explanation must remain speculative.

### Experiment 3

This experiment provided the final test of the effects of manipulating online control on motor imagery and physical action. Here, participants engaged in either the standard reaching and placing task, or a mimed version of the task using a faux disc. The *Mime* condition would be presumed to impair normal online control processes, as these are argued to rely on physical interaction with the target in the environment (Goodale et al., 1994).

The predictions were similar to those in the previous experiments: If the Motor Cognitive model is correct, we would expect to see an effect of *Mime* whereby only physical actions would be affected by whether the movement was real or mimed, resulting in an *Action* by *Mime* interaction. Further, we would also expect the typical main effect of *Action*, whereby motor imagery would take longer than physical actions. Conversely, if the Functional Equivalence view is correct, only the main effect of *Mime* should be observed.

## Methods

### Participants

We enrolled 60 participants (43 women, 17 men, tested age range 18-43 years old) from Royal Holloway students and staff, 30 participants in each group. Four participants were replaced because they failed to follow the instructions. Inclusion criteria and Covid-19 safety precautions were the same as in Experiments 1 and 2. Sample size and inclusion/exclusion criteria were determined *a priori* and were pre-registered prior to data collection alongside study procedures and data processing and analyses (https://osf.io/ws2zn). The full dataset and scripts are available to download at the following link: https://osf.io/a9tzf/. Data collection took place from March to August 2022.

### Experimental setup and procedure

Setup, starting position and number of trials were the same as in Experiments 1, 1a and 2 for one of the conditions (“Control”; see left panel of Figure 5). For the second condition (“Mime”), in place of the disc was a paper circle faux disc fastened to the table using tape, in the usual location of the disc. The real disc was placed 12 cm to the right of the faux disc (centre to centre, i.e. space of 8 cm between the paper and the real discs, see right panel of Figure 5) so that participants could see it clearly during their tasks.

**Figure 4.**
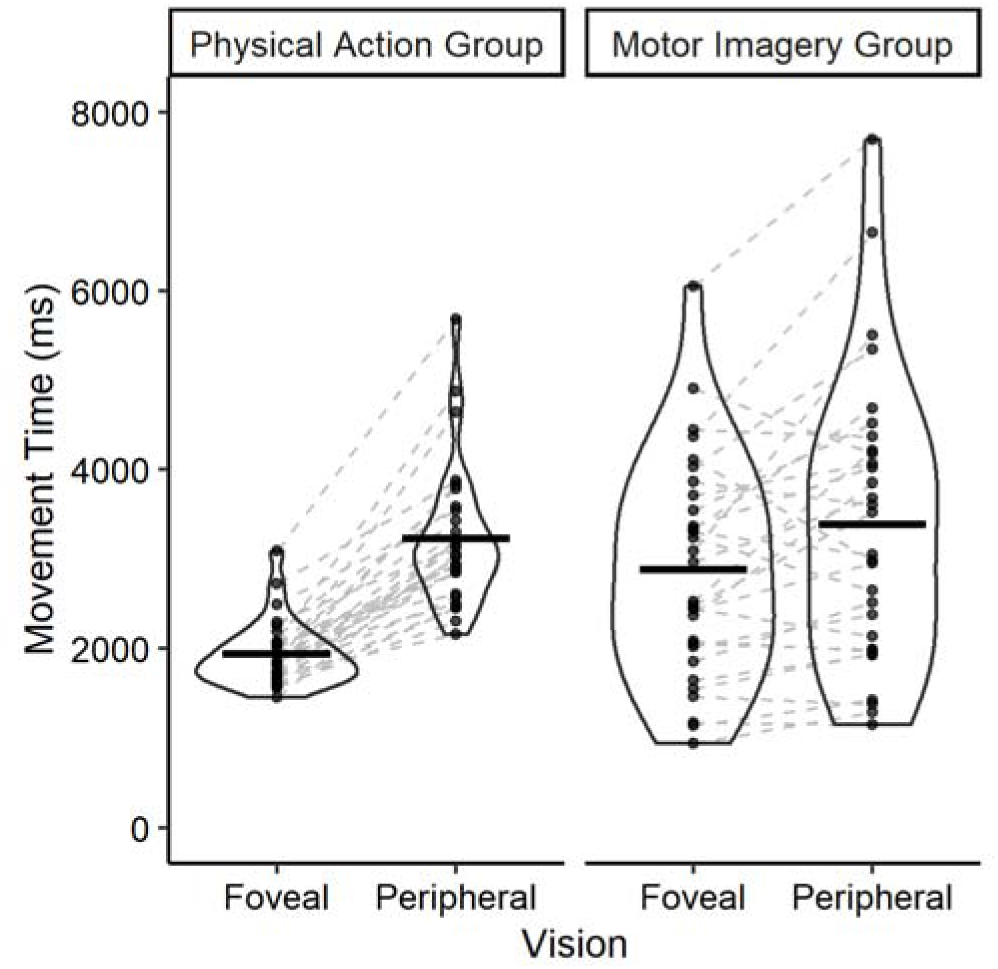
Effect of varying the quality of online control (Foveal vs. Peripheral) on movement times in the physical action and motor imagery groups. Conventions as in Figure 2.

**Figure 5.**
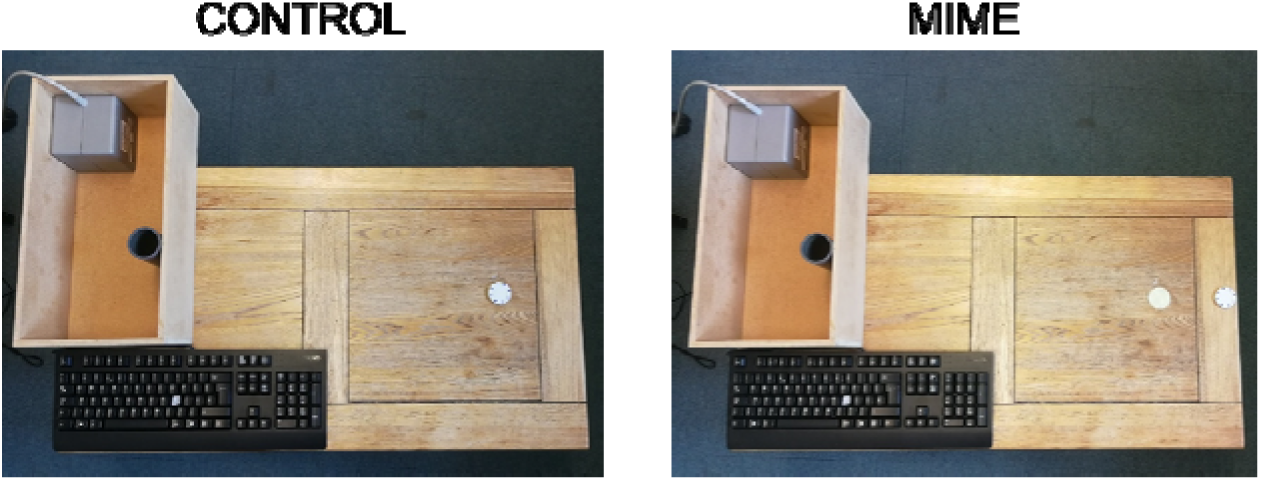
Experimental setup for the Control and the Mime conditions. In the mime condition, the disc was moved 12 cm to the right (centre to centre) and replaced by an identically sized circle of white paper.

**Figure 6.**
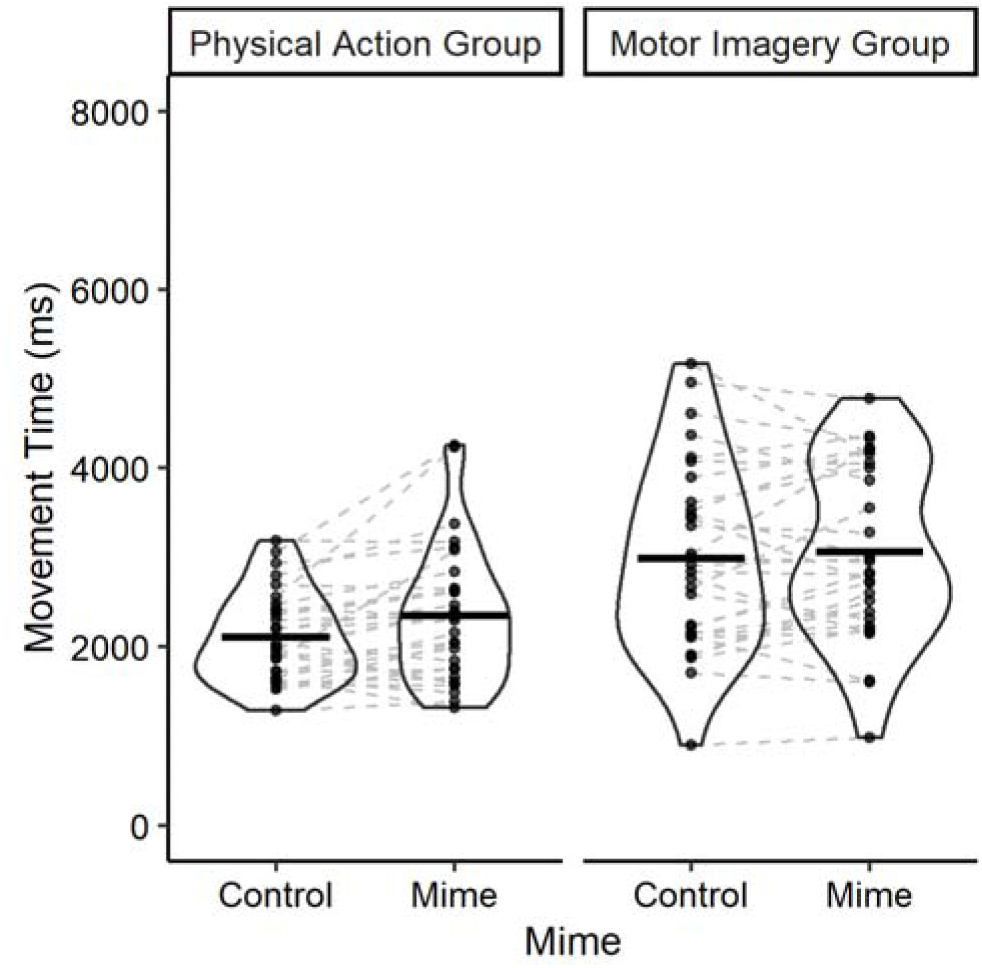
Effect of varying the quality of online control (Control vs. Mime) on movement times in the physical action and motor imagery groups. Conventions as in Figure 2.

One group of participants performed the physical action under normal conditions (Control trials) or by pantomiming the action (Mime trials). This latter condition consisted of performing the same movement but using the faux disc: participants reached towards the faux disc and moved their (empty) fingers towards the cylinder. The second group imagined reaching and placing the faux disc in the same two conditions. The experimenter continuously monitored the participants during each trial to ensure they followed instructions. Out of 1440 trials in total, 53 (3.7%) were excluded and not repeated due to participants’ mistakes such as movement during motor imagery, participants forgetting to press the button (Motor Imagery Control: 5/360; Motor Imagery Mime: 8/360; Physical Action Control: 25/360; Physical Action Mime: 15/360).

### Data processing

Data processing was performed exactly as in Experiments 1, 1a and 2. Additionally to the non-analysed trials, in the keypress data, 3.2% of the trials had RT or MT outside ± 2 IQR (Interquartile Range) for each combination of block and were removed (Motor Imagery Control: 0.6%; Motor Imagery Mime: 0.9%; Physical Action Control: 0.4%; Physical Action Mime: 1.2%). In the kinematic data, the number of trials removed was 2.8% (Control: 1.0%; Mime: 1.8%).

### Analyses and statistics

Analyses were the same as in Experiments 1 and 2, except that the *Mime* condition replaced the *Vision* condition as the within-subjects variable.

## Results

### Kinematic indices

Table 4 shows the effects of *Mime* on the kinematic indices in the physical reaching and placing task. There was strong evidence that performing the mimed action increased overall movement times (λ_adj_ = 63, *dz* = .64). The evidence for an increase in time spent in the deceleration phase during the placing phase was more moderate (λ_adj_ = 5.9, *dz* = .45), whereas evidence for an increase in the number of online adjustments during the placing phase was strong (λ_adj_ = 14.2, *dz* = .52). However, neither time spent in deceleration during the reaching phase (λ_adj_ = 1.3, *dz* = .30), nor the number of online adjustments during the reaching phase were noticeably different between conditions (λ_adj_ = 1.9, *dz* = .34); these particular results were at odds with our predictions of an effect of *Mime* on online control. Overall, these results suggest that our manipulation had much more modest effects on online control than were present in Experiments 1, 1a, and 2. As before, there was very little difference between movement times indicated by the kinematics versus the keypress, suggesting that participants were accurate in indicating the beginning and end of the movement. This is in accordance with an additional analysis performed on kinematic MT for the physical action group vs. keypress MT for the motor imagery group which rendered the same results as in the next section (see Supplementary Information).

**Table 4.** Effect of performing a movement on kinematics in the Control and Mime conditions.

| | Control (mean ± SD) | Mime (mean ± SD) |
| --- | --- | --- |
| Total movement time (ms) | 2072 ± 492 | 2355 ± 742 |
| **Reaching phase** |  |  |
| Deceleration Time (ms) | 679 ± 185 | 749 ± 327 |
| Number of online adjustments | 1.3 ± 1.0 | 1.8 ± 1.9 |
| **Placing phase** |  |  |
| Deceleration Time (ms) | 795 ± 192 | 879 ± 287 |
| Number of online adjustments | 1.7 ± 1.1 | 2.2 ± 1.1 |

### Keypress movement times

Figure 5 shows the effects of *Action* and *Mime* on keypress movement times. The best-fitting model of keypress MT included the main effects of *Action* and *Mime*, but no interaction. A statistical model including an effect of *Action* fit the data much better than a null model (λ_adj_ = 449, *f*² = 1.58), with longer movement times in the motor imagery group than in the physical action group. Adding a main effect of *Mime* moderately improved the fit (λ_adj_ = 7.5, *f*² = .37), with longer movement times in the mime condition than in the control condition. Contrary to the prediction of the Motor-Cognitive model, adding the interaction *Action* × *Mime* did not improve the fit (λ_adj_ = 0.97, *f*² = .09). Nonetheless, despite the lack of evidence for the predicted interaction, the data overall were again more consistent with the Motor-Cognitive model than with the Functional Equivalence view: A full model including the effects of *Action, Mime,* and their interaction fit the data much better than a model including only a main effect of *Mime,* (λ_adj_= 433).

## Discussion

The kinematic data in Experiment 3 was less compelling than in the previous two experiments. Although there was moderate-to-good evidence that three of the five kinematic indices were affected by the Mime condition, for the other two indices the evidence was inconclusive. As such, it seems fair to conclude that the *Mime* manipulation had a smaller effect on online control than either the no vision or the peripheral vision manipulations in the first three experiments.

Although imagined movement times were longer than physical movement times, the predicted *Action* × *Mime* interaction did not arise. Overall, however, the pattern of results was more consistent with the Motor Cognitive model than the Functional Equivalence view, due to the large main effect of *Action*.

A possible explanation for the lack of evidence for an *Action* × *Mime* interaction was that the effect of *Mime* was small compared to the comparable manipulations of online control used in the other two experiments. As a consequence, although the pattern of means was consistent with *Mime* having a greater effect on physical actions than on motor imagery, with an effect of 240 msec on physical action compared to 70 msec for motor imagery, the statistical evidence for the interaction was lacking. It is possible the predicted interaction was present but much smaller than we had predicted and may require a larger sample size to be reliably detected.

### General Discussion

Keypress movement time data in all four experiments supported the Motor Cognitive model over the Functional Equivalence model. Critically, participants who imagined the movements were consistently less affected by the manipulations that impacted online control, in line with the predictions of the Motor-Cognitive model. However, physical actions were affected by these same variables (although the evidence for this in Experiment 3 was modest). Overall, these results are incompatible with the view of physical action and motor imagery being functionally equivalent.

Whereas the key interactions between *Action* and online control were highly evident in Experiments 1, 1a and 2, in Experiment 3 the smaller and less consistent effects of our manipulation on online control made it hard to discern if an interaction between *Action* and *Mime* was present, resulting in a lack of clear evidence either favouring or refuting an interaction. We conclude from this that the impact on online control in this experiment was insufficient to allow clear evidence of the interaction, although the effect of *Action,* uniquely predicted by the Motor-Cognitive model, was still present. It may be the case that online control is not as severely disrupted by miming a movement as has been previously argued (Goodale et al., 1994).

Results from Experiment 2 were not entirely consistent with a recent study which also found that the inability to perform eye movements (i.e. peripheral trials) lengthened motor imagery times (Pathak et al., 2023), but which observed the opposite pattern for the physical action. In that study, physical action was shorter in the condition that did not allow eye movement, while we observed longer movement times in this condition. At first glance, the opposite pattern observed in Pathak et al. (2023) contrasted with both the predictions of the Motor Cognitive model and the results of Experiment 2. However, we suspect this is likely due to experimental differences. Whereas both that study and the present study involved participants actively inhibiting eye movements in one condition, the two studies otherwise differed in significant ways. Specifically, whereas the Fitt’s type task used by Pathak et al. maximised the use of feedforward strategies, the present study focussed on a change in the quality of online control, which in turn affected movement times of the physical action. Regarding motor imagery, we suspect the effects may be due to difficulties in imagining movements while maintaining fixation. More generally, it would be interesting to test the effects of restricting eye movements on motor imagery independently of other instructions, to see if it disrupts the ability of participants to carry out motor imagery.

The present study represents part of a larger experimental dissociation between motor imagery and physical action. In previous work, we demonstrated that numerous variables, including precision, cognitive load, and disruption of the dorsolateral prefrontal cortex with TMS, all impacted the timing of motor imagery much more than the corresponding physical actions (Glover et al., 2020; Glover & Baran, 2017; Martel & Glover, 2023). Here, we have shown the opposite pattern: variables that have large effects on physical action may have much smaller effects on motor imagery. Although either pattern of effects alone might be argued to arise as an artefact of the reaching and placing paradigm we are using, it seems *prima facie* extremely unlikely for such an artefact to produce opposing patterns of results depending on the manipulations employed. Further, our control analyses showed that the keypress movement times for physical actions closely corresponded to the kinematic measures of same, again refuting the idea of an artefact producing the results. We thus conclude that not only are there clear differences between motor imagery and physical action, but the processes are largely if not wholly independent during execution. The presence of smaller, but present, effects on motor imagery version physical action here, and the opposite pattern in previous studies (Glover et al., 2020; Glover & Baran, 2017; Martel & Glover, 2023), are wholly consistent with this view, as motor imagery and physical action are argued to share a common planning stage. Thus, one would not expect a complete dissociation of the effects of different variables on the two behaviours.

### Other Accounts of Motor Imagery

Following calls for further theoretical development in motor imagery research (Dietrich, 2008; Guillot et al., 2012; Moran et al., 2012; O’Shea & Moran, 2017), several other promising theoretical frameworks have been put forth (e.g., Bach et al., 2022; Frank et al., 2023; Grush, 2004; Hurst & Boe, 2022; Rieger et al., 2017). We will discuss three of these models in turn, and how they might attempt to explain the results of the present study (Frank et al., 2024; Grush, 2004; Rieger et al., 2017).

First, in his Motor Emulation Model, Grush (2004) argued that motor imagery is the conscious experience of the forward model when the planned movement is inhibited. By emphasizing the use of a forward model, this view differs from the Functional Equivalence view in terms of how motor commands are executed. Yet both contend that motor imagery arises from internal motor processes without any influence from other cognitive inputs, As such, the Grush (2004) model might be expected to make broadly similar predictions to the Functional Equivalence view, with some notable exceptions such as predicting that motor imagery can result in sensory attenuation (Kilteni et al., 2018).

A champion of the Motor Emulation model might try to explain the results of the present study as being a consequence of differing amounts of feedback influencing motor imagery. For example, removing vision might cause motor imagery times to lengthen because the emulator anticipates the lack of visual feedback and requires more time to use (simulated) proprioceptive feedback to execute the motor image. If this were the case, however, one would not expect there to be an interaction between motor imagery and vision as was observed here, as the forward model should experience the same difficulties whether the action was carried out physically or imagined. Moreover, the Grush (2004) view would have difficulty explaining our previous findings, such as the effects of interference tasks on motor imagery that are much larger than on physical action (Glover et al., 2020; Glover & Baran, 2017; Martel & Glover, 2023), or effects of disrupting the DLPFC on motor imagery but not physical action (Martel & Glover, 2023), given that it does not consider the influence of cognitive inputs on motor imagery. In sum, it is difficult to account for these recent findings by positing that motor imagery relies solely on a forward model-based emulator.

Second, the Inhibition Hypothesis of Rieger and colleagues (Rieger et al., 2017) agrees with the Motor Cognitive view that motor imagery is not simply a byproduct of internal motor processes that occur during motor planning. Rather, they contend that motor imagery incorporates inhibition processes that necessarily slow its performance relative to physical action. On this account, motor imagery might be expected to consistently take longer than the corresponding physical action. This hypothesis has some support from studies showing that, for short-duration movements at least, motor imagery tends to have greater movement times than physical action (Guillot & Collet, 2005). There are two issues the Inhibition hypothesis would have in explaining the present study, however. For one, whereas we did observe longer movement times for motor imagery than for physical action in many of the conditions tested here, the opposite was found in the No Vision conditions of Experiments 1 and 1a, wherein physical actions took longer than motor imagery. It seems difficult to reconcile this with the idea that inhibition of movements in motor imagery ought to result in longer movement times in motor imagery than the corresponding physical action. For another, even if one were able to explain these contradictory results in terms of inhibition, inhibition effects in cognition generally are on the order of several tens of milliseconds (e.g., the Stroop effect, MacLeod, 1991), whereas the differences observed here and in our previous studies are on the order of hundreds of milliseconds (Glover et al., 2020; Glover & Baran, 2017; Martel & Glover, 2023). Thus, although we can accept the notion that inhibition, as an executive function, may contribute to slowing motor imagery under certain conditions, it cannot be the only explanation.

Third is the Perceptual-Cognitive Scaffolding Model of Frank et al. (2024). According to this view, motor imagery is a simulation of action, effectively predicting the perceptual consequences of the inhibited movement in a similar way to that argued for physical action in the Theory of Event Coding (Hommel et al., 2001). By elaborating on the construction of an increasingly sophisticated perception-action scaffolding, motor imagery can facilitate motor learning (Frank et al., 2024). This framework does well at explaining how motor imagery training enhances physical performance (Simonsmeier et al., 2021; Toth et al., 2020). Although this theoretical framework is largely orthogonal to the aims of the present study, the Perceptual-Cognitive Scaffolding Model could be used to explain the results of Experiment 1a, wherein practice with motor imagery did not affect the timing of physical actions. To account for this, a proponent of the Frank et al. (2024) framework would likely contend that either the amount of training in motor imagery was insufficient to facilitate motor learning, or that the movement itself was too simple to enable the kind of complex perception-action scaffolding that benefits learning. The Perceptual-Cognitive Scaffolding Model, while not directly tested here, is nonetheless consistent with our findings.

### Limitations

At least two critiques of the present study may be offered. First, the presence/absence of eye movements or surreptitious opening/closing of the eyes were not controlled for using eye-tracking or vision-blocking technology and it is not impossible that the experimenter missed trials that should have been excluded. However, the fact that both kinematics and keypress data showed longer movement times in the Physical Action, no vision/peripheral vision conditions of Experiments 1, 1a, and 2, suggests that the participants conformed with instructions, as the manipulations had the predicted effects on online control.

A second limitation is that three of the four experiments reported here used a between-participants design. This precluded the possibility of observing the effects of performing one type of behaviour on later performance of the other. However, we previously observed that such practice did not materially alter the effects germane to the predictions of the Motor Cognitive model (Martel & Glover, 2023), and the same outcome was observed here in Experiment 1a. Regardless of whether some participants had prior experience with the physical version of the task, the predicted interaction between action and vision remained, motor imagery being less affected by the manipulation of online control than physical actions. We thus conclude that regardless of prior experience with a task, motor imagery is unable to incorporate online control into its performance, but instead uses executive functions to monitor and elaborate the motor image during execution.

## Acknowledgments

This work was supported by the Leverhulme Trust (grant number RPG-2020-225).

## Constraints on generality statement

The present study supported the Motor-Cognitive Model with four different experiments testing healthy, right-handed, and generally youthful university students. We have no reason to believe that the results depend on other characteristics of the participants, materials, or context.

