## Supplementary Material for "Online versus Cognitive Control: A Dividing Line between Physical Action and Motor Imagery"

**Marie Martel & Scott Glover**

### 1. Experiment 1

#### a. Additional analysis on MT

As an additional check of our results, we performed a second (non-preregistered) analysis of the data using the keypress MT for the motor imagery group and the kinematic MT for the physical action group (Figure S1). If the results are similar, then we can consider with high confidence that keypresses served as a good proxy for movement onset/offset.

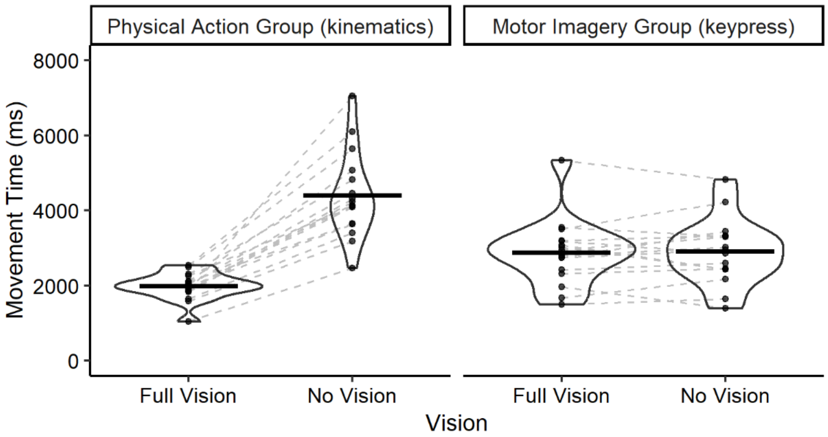


**Figure S1. Effect of varying vision (Full Vision vs. No Vision) on movement times in the physical action and motor imagery groups using kinematics and keypress MT respectively.** Individual points represent the mean score for single participants, dashed grey lines pair the means of individual participants together, and horizontal lines represent the group means. Enclosures represent the distribution of the presented data, with wider horizontal sections indicating a larger number of observations.

Similar to the analysis in the main manuscript, the best-fitting model of the data included the effect of *Vision* and the *Vision* × *Action* interaction. A statistical model including only an effect of *Action* did not fit the data noticeably better than a null model ($\lambda_{adj} = 2.6$), however, adding the effect of *Vision* ($\lambda_{adj} > 1000$) improved the fit greatly, and adding the interaction *Action* × *Vision* ($\lambda_{adj} > 1000$) further improved the fit, reflecting increased movement times in the no vision condition but only for the physical action group (Figure S1).

This analysis shows that the keypresses accurately indexed movement times in the physical action group, supporting the robustness of our experimental design (compare the left panel of Figure 2 from the main manuscript to Figure S1 below).

#### b. Keypress Reaction Times

Neither the Motor-Cognitive model nor the Functional Equivalence view make any specific predictions regarding the RT. We thus conducted an exploratory analysis of the data, which is presented in Figure S2. The best-fitting model included only an effect of *Action*: participants took longer to initiate an imagined movement compared to the physical action. A statistical model including an effect of *Action* fit the data much better than a null model ($\lambda_{adj} > 1000$), with longer reaction times for the Motor Imagery group. Adding a main effect of *Vision* did not improve the fit ($\lambda_{adj} = .67$), nor did adding an interaction *Action* × *Vision* ($\lambda_{adj} = .48$).

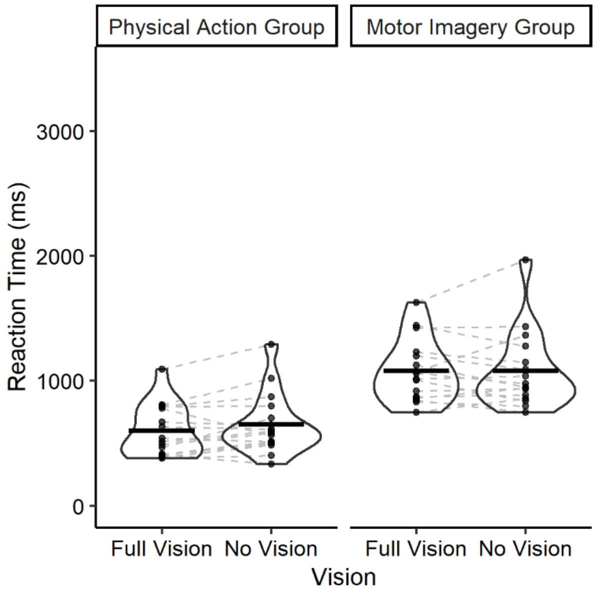


**Figure S2. Effect of varying vision (Full Vision vs. No Vision) on reaction times in the physical action and motor imagery groups.** Individual points represent the mean score for single participants, dashed grey lines pair the means of individual participants together, and horizontal lines represent the group means. Enclosures represent the distribution of the presented data, with wider horizontal sections indicating a larger number of observations.

#### c. First check for keypress integrity: Individual kinematic correlations

*Methods.* As a methodological control, we initially measured the correspondence between button-pressing and actual reaction times and movement times measured using the Polhemus motion tracking system. Keypress RT and MT were calculated as defined in the main manuscript. Kinematic RT was defined as the time between the beep and the beginning of the grasping component, that is the time when the hand first reaches a velocity of 5cm/s after initiating the movement. MT was defined in the three following ways: 1) the end of the placing movement based on a velocity criterion of when the velocity had reached its minimum or had fallen below 5cm/s after the grasping component of the movement, and before the participants had moved back to the starting position; 2) the end of the movement based on a consistent increase in grip aperture occurring after the end of the grasping movement (indicating the beginning of the release of the disc from the hand); and 3) the end of the movement based on the maximum grip aperture occurring after the end of the grasping movement (indicating the full release of the disc). The RT and MT as measured by these methods were then compared to RT and MT data from the keypresses, with correlations measured to index the efficacy of these kinematic markers in predicting the timing of the button. Data from a previous experiment (Martel & Glover, 2023) showed that this was a broadly effective way of testing the integrity of the participants' button–press times. We applied Pearson correlations for each participant separately for each *Vision* condition.

*Results.* Moderate correlations were observed between the keypressing indicating the start of the movement and the start of the movement based on the kinematic measure (RT), as well as between the keypressing indicating the end of the movement and the time of the end of the movement given by our three different kinematics markers (MTvel, MTinc and MTmga; Figure S3). Correlations with the velocity-based end of movement were the highest. The other criteria were less optimal in determining the end of the movement, probably because as compared to the previous study in which we used these markers (Martel & Glover, 2023), the movement was different so people did not need to open their fingers during the movement as they released the disc only at the very end of the movement.

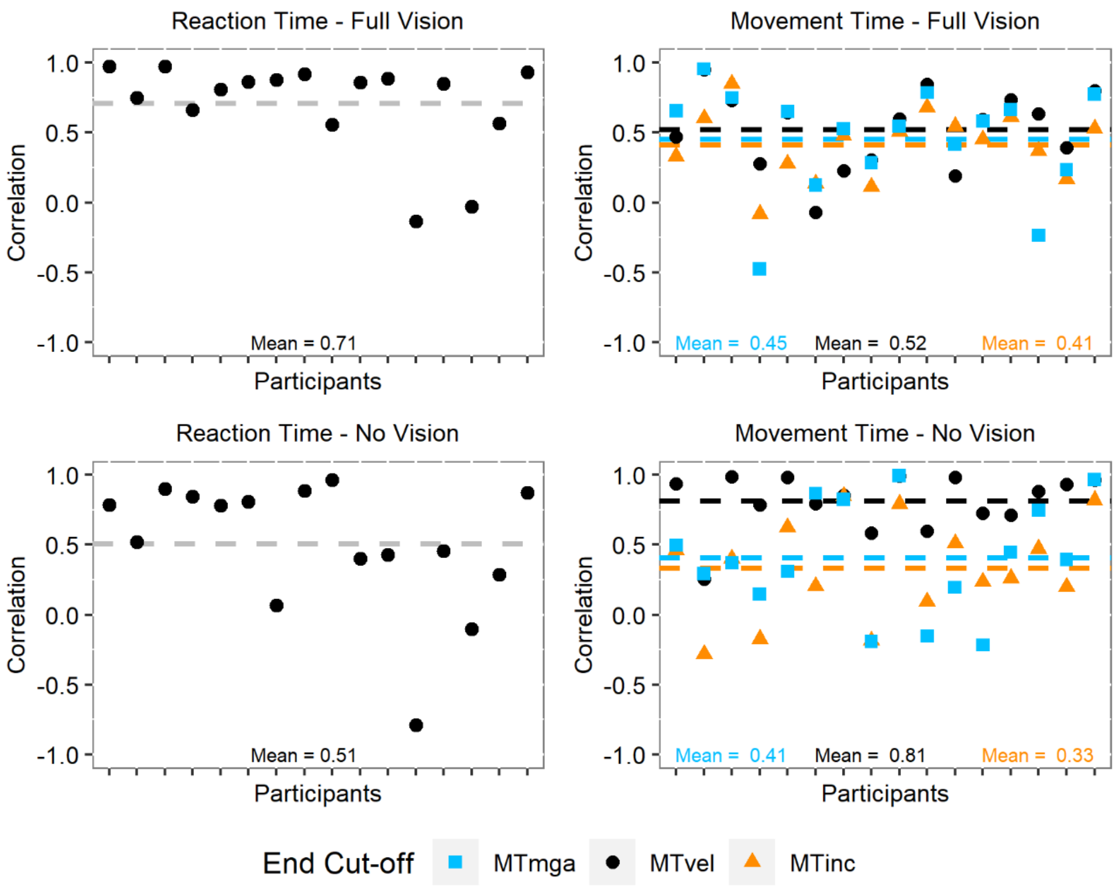


**Figure S3. Pearson correlation values between the keypress data and the kinematics parameters in Experiment 1.** Symbols are individual correlation values for the reaction times (left panel) or the movement times (right panel). In the right panel, MT values on the same column belong to the same participant. Dashed lines indicate the mean correlation for each marker. MT = Movement Time, defined by 3 different criteria: lowest velocity (vel), increased grip aperture (inc) or maximum grip aperture (mga) during the second part of the movement.

#### d. Two additional checks for keypress integrity based on group means

It is notable from Figure S3 that individual correlations between keypress and kinematic MTs are highly variable. Although this might suggest that keypress times were inaccurate indexes of movement start and stop times in some participants, on reflection we believe a more likely explanation is that this simply reflects the lack of variability in movement times within blocks, resulting in it being difficult to capture the effects using correlations on an individual level.

A better approach would be to measure the correlation between the mean MTs for all participants across all trials with the comparable mean kinematic MTs. This analysis shows a very high correlation, $r^2$ =

0.93 for the Full Vision and $r^2 = 0.98$ for the No Vision condition, showing a very strong overall correspondence between the two measures (see Figure S4).

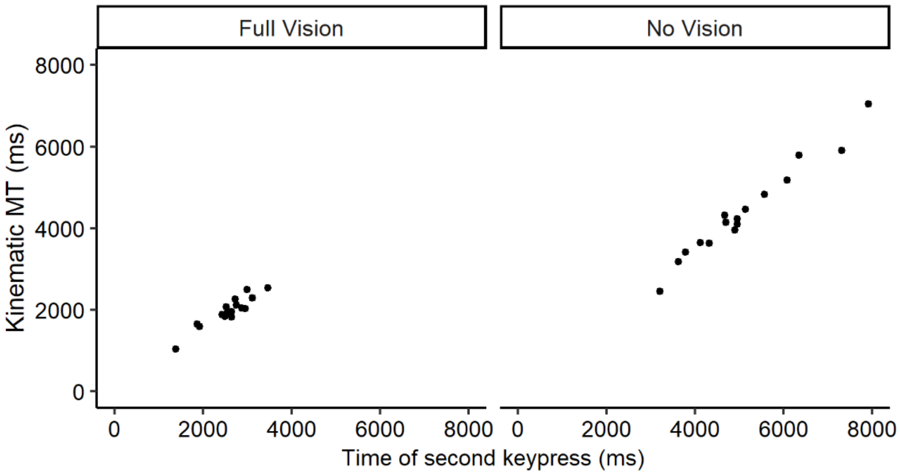


**Figure S4. Correlations between kinematic and keypress MT for each participant across all trials, in each condition.**

A direct comparison of means across the different measures confirms this view. As can be seen below (Figure S5), there was no systematic bias in keypressing, with nearly identical mean movement times across the different conditions.

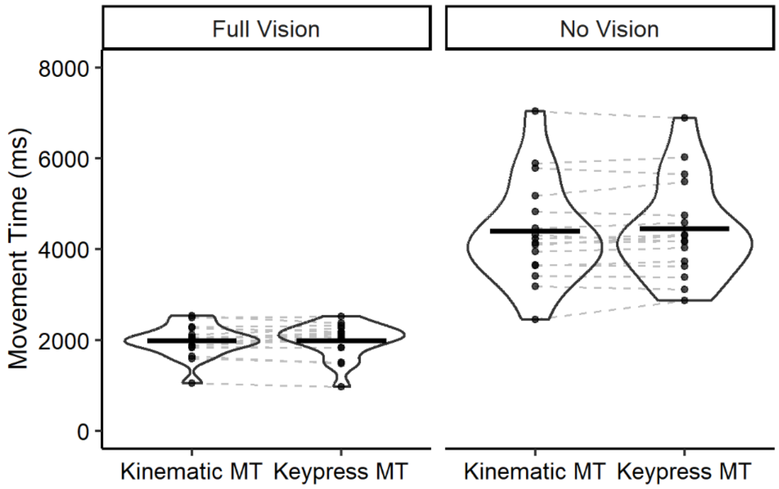


**Figure S5. Comparison between kinematic and keypress movement times in the physical action group.** Individual points represent the mean MT for single participants, dashed grey lines pair the means of individual participants together, and horizontal lines represent the group means. Enclosures represent the distribution of the presented data, with wider horizontal sections indicating a larger number of observations.

In short, we can be confident that keypress movement times accurately reflected the movement times measured through kinematics, and that there were no artefactual effects present in the analyses presented in the main manuscript.

### 2. Experiment 1a

#### a. Additional analysis on MT

As an additional check of our results, we performed the same extra analyses as for Experiment 1.

Similar to the analysis in the main manuscript, the best-fitting model using kinematic MT instead of keypress MT was a full model including the effects of *Action*, *Vision* and their interaction (Figure S6), with an additional effect of *Order*. A statistical model including only an effect of *Action* fit the data much better than a null model ($\lambda_{adj} > 1000$) with longer movement times in the physical action than in the motor imagery group. Adding *Vision* to the model further improved the fit ($\lambda_{adj} > 1000$), with longer movement times in the absence of Vision. Adding the interaction *Action* × *Vision* vastly improved the fit ($\lambda_{adj} > 1000$): the effect of *Vision* was greater in the Physical Action group than in the Motor Imagery group. Finally, adding the effect of *Order* moderately increased the fit ($\lambda_{adj} = 7.7$) with overall longer MT when participants started with Motor Imagery.

This analysis shows that the participants pressed the key precisely in the physical action group, supporting our experimental design (compare the left panel of Figure 3 from the main manuscript to Figure S6 below).

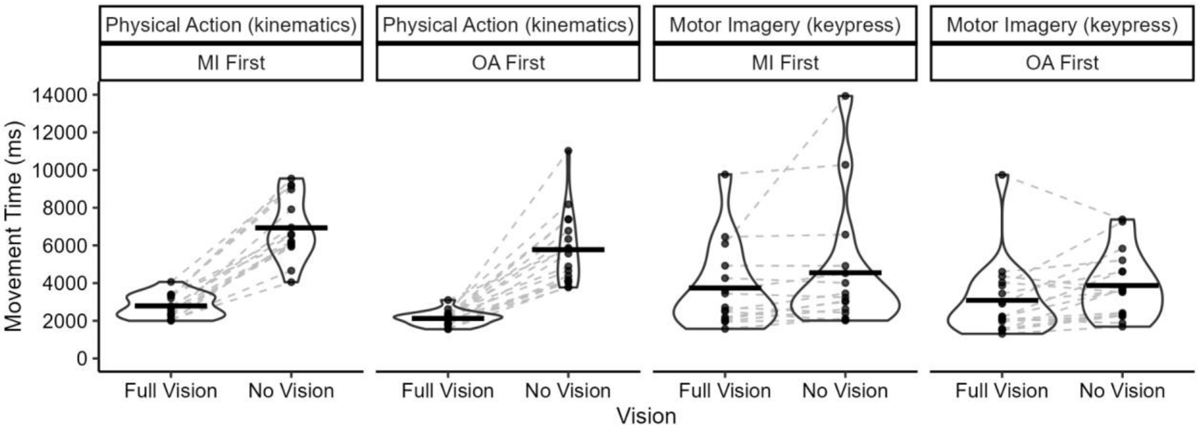


**Figure S6. Effect of varying vision (Full Vision vs. No Vision) on movement times in the physical action and motor imagery conditions using kinematics and keypress MT respectively, depending on the order of the tasks.** Individual points represent the mean score for single participants, dashed grey lines pair the means of individual participants together, and horizontal lines represent the group means. Enclosures represent the distribution of the presented data, with wider horizontal sections indicating a larger number of observations.

#### b. Keypress Reaction times

Neither the Motor-Cognitive model nor the Functional Equivalence view make any specific predictions regarding the RT. We thus conducted an exploratory analysis of the data, which is presented in Figure S7. The best-fitting model included main effects of *Action* and *Vision* and an interaction between *Order* and *Action*. A statistical model including an effect of *Action* fit the data much better than a null model ($\lambda_{adj} > 1000$), indicating that participants took longer to initiate an imagined movement compared to the physical action, so did a model including a main effect of Vision ($\lambda_{adj} > 1000$), showing that participants too longer in absence of action. Adding the interaction *Order* × *Action* further improved the fit ($\lambda_{adj} = 416$), suggesting that RT were similar for the Imagery condition, irrespective of the Order of the conditions, while participants initiated their movement faster when they performed the physical action if they started with Physical Action (instead of Motor Imagery).

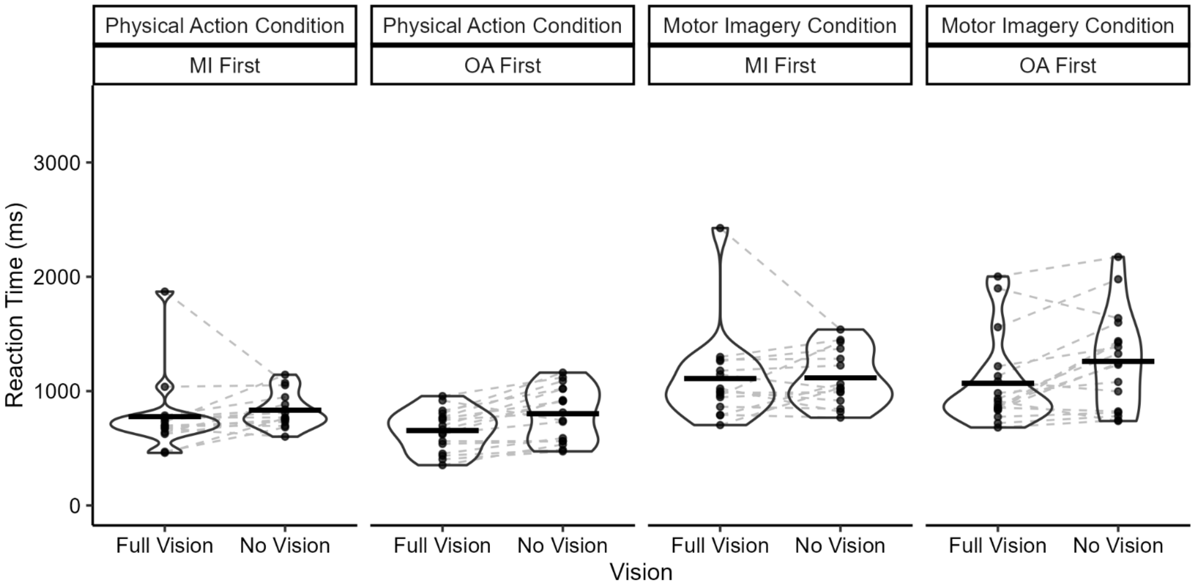


**Figure S7. Effect of varying the quality of online control (Full Vision vs. No Vision) on reaction times in the physical action and motor imagery conditions, depending on the order of the task.** Conventions as in Figure S2.

#### c. First check for keypress integrity: Individual kinematic correlations

*Methods.* We again measured the correspondence between button-pressing and actual reaction times and movement times measured using the Polhemus motion tracking system, using the same criteria as in Experiment 1. We applied Pearson correlations for each participant and each *Vision* condition.

*Results.* Moderate correlations were observed between the keypressing indicating the start of the movement and the start of the movement based on the kinematic measure (RT), as well as between the keypressing indicating the end of the movement and the time of the end of the movement given by our three different kinematics measures (MTvel, MTinc and MTmga; Figure S8). Correlations with the velocity-based end of movement were again the highest.

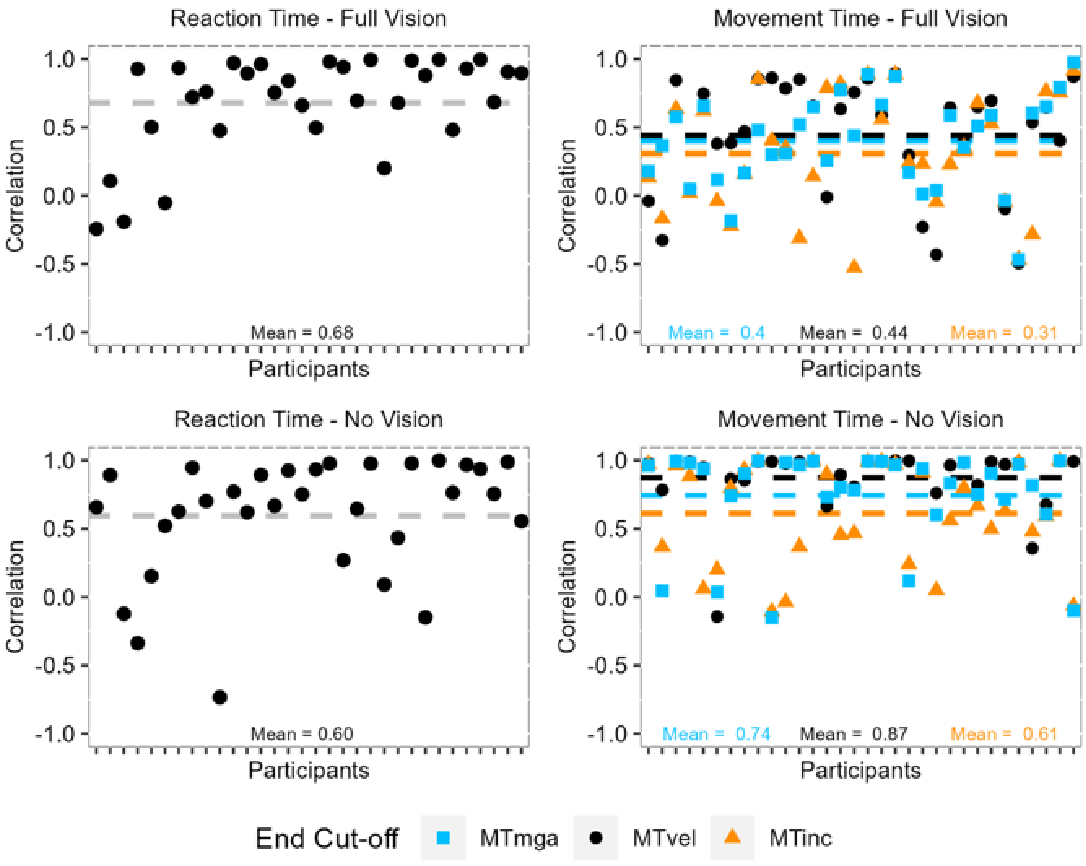


**Figure S8 Pearson correlation values between the keypress data and the kinematics parameters in Experiment 1a.** Symbols are individual correlation values for the reaction times (left panel) or the movement times (right panel). In the right panel, MT values on the same column belong to the same participant. Dashed lines indicate the mean correlation for each marker. MT = Movement Time, defined by 3 different criteria: lowest velocity (vel), increased grip aperture (inc) or maximum grip aperture (mga) during the second part of the movement.

#### d. Two additional checks for keypress integrity based on group means

It is again notable that individual correlations between keypress and kinematic MTs were highly variable (Figure S8). Although this might suggest that keypress times were inaccurate indices of movement start and stop times in some participants, on reflection we believe a more likely explanation is that this simply reflects the lack of variability in movement times within blocks, resulting in it being difficult to capture the effects using correlations on an individual level.

As before, a better approach would be to measure the correlation between the mean MTs for all participants across all trials with the comparable mean kinematic MTs. This analysis shows a very high correlation, $r^2 = 0.84$ in the Full Vision condition and $r^2 = 0.97$ in the No Vision condition, showing a very strong overall correspondence between the two measures (see Figure S14).

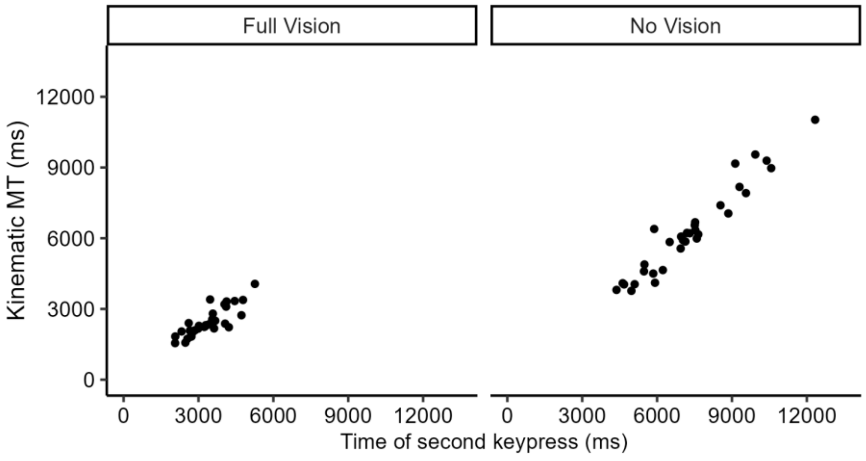


**Figure S9. Correlations between kinematic and keypress MT for each participant across all trials, in each condition.**

A direct comparison of means across the different measures confirms this view, with a good match in both conditions between the keypress and the kinematic MT.

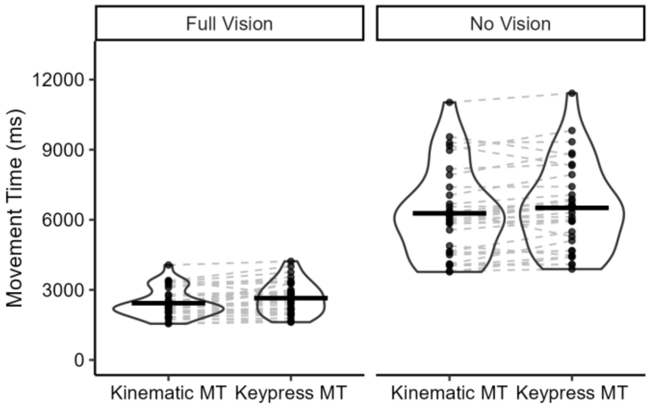


**Figure S10. Comparison between kinematic and keypress movement times in the physical action group.** Conventions as in Figure S4.

### 3. Experiment 2

#### a. Additional analysis on MT

As an additional check of our results, we performed the same extra analyses as for Experiment 1.

Similar to the analysis in the main manuscript, the best-fitting model using kinematic MT instead of keypress MT was a full model including the effects of *Action*, *Vision* and their interaction (Figure S11). A stastistical model including only an effect of *Action* fit the data much better than a null model ($\lambda_{adj}$ >

1000) with longer movement times in the motor imagery group than in the physical action group. Adding *Vision* to the model further improved the fit ($\lambda_{adj}$ > 1000), with longer movement times in the Peripheral condition than in the Foveal condition, a finding in the opposite direction to what we predicted. Finally, adding the interaction Action × Vision vastly improved the fit ($\lambda_{adj}$ = 22): the effect of Vision was greater in the physical action group than in the motor imagery group.

This analysis shows that the participants pressed the key precisely in the physical action group, supporting our experimental design (compare the left panel of Figure 4 from the main manuscript to Figure S11 below).

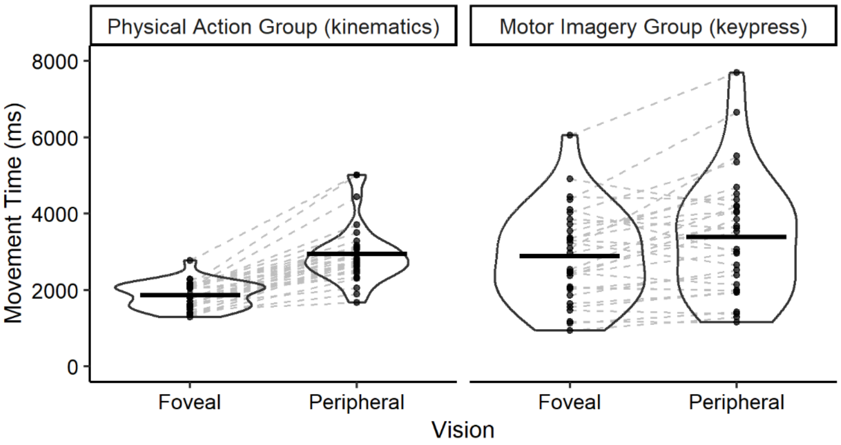


**Figure S11. Effect of varying the quality of online control (Foveal vs. Peripheral) on movement times in the physical action and motor imagery groups.** Conventions as in Figure S1.

#### b. Keypress Reaction times

Neither the Motor-Cognitive model nor the Functional Equivalence view make any specific predictions regarding the RT. We thus conducted an exploratory analysis on the data, these are presented in Figure S12. The best fitting model included main effects of *Action* and *Vision*: participants took longer to initiate an imagined movement compared to the physical action and took longer in the peripheral condition than the foveal one. A statistical model including an effect of *Action* fit the data much better than a null model ($\lambda_{adj}$ > 1000), so did a model including a main effect of *Vision* ($\lambda_{adj}$ > 1000). Adding the interaction *Action* × *Vision* marginally improved the fit ($\lambda_{adj}$ = 2.6), suggesting that the increase in the peripheral relative to foveal condition was possibly greater in the motor imagery group than in the physical action group.

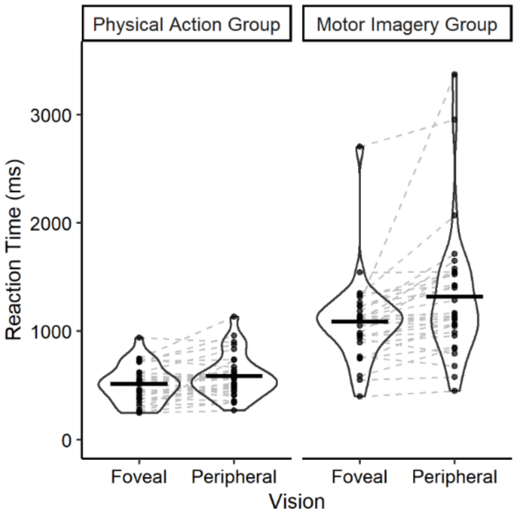


**Figure S12. Effect of varying the quality of online control (Foveal vs. Peripheral) on reaction times in the physical action and motor imagery groups.** Conventions as in Figure S2.

#### c. First check for keypress integrity: Individual kinematic correlations

*Methods.* We again measured the correspondence between button-pressing and actual reaction times and movement times measured using the Polhemus motion tracking system, using the same criteria as in Experiment 1. We applied Pearson correlations for each participant and each *Vision* condition.

*Results.* Moderate correlations were observed between the keypressing indicating the start of the movement and the start of the movement based on the kinematic measure (RT), as well as between the keypressing indicating the end of the movement and the time of the end of the movement given by our three different kinematics measures (MTvel, MTinc and MTmga; Figure S13). Correlations with the velocity-based end of movement were again the highest.

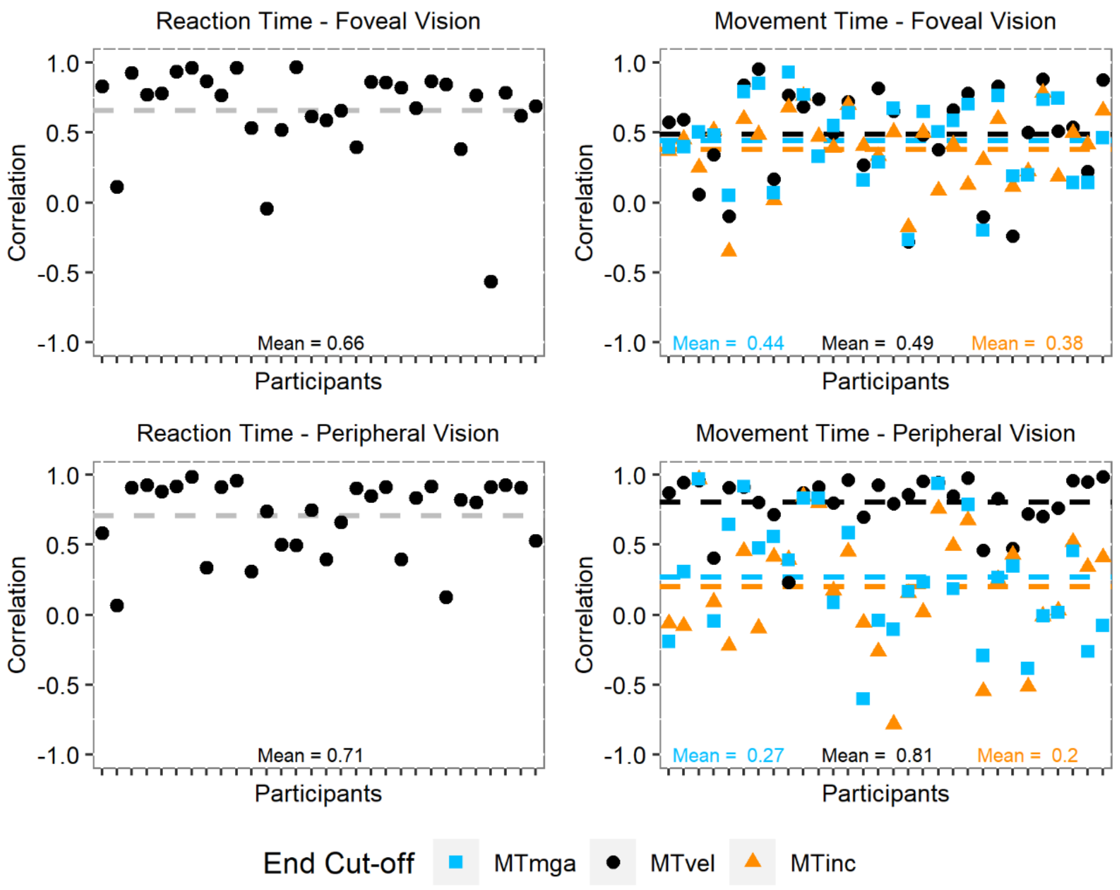


**Figure S13. Pearson correlation values between the keypress data and the kinematics parameters in Experiment 2.** Symbols are individual correlation values for the reaction times (left panel) or the movement times (right panel). In the right panel, MT values on the same column belong to the same participant. Dashed lines indicate the mean correlation for each marker. MT = Movement Time, defined by 3 different criteria: lowest velocity (vel), increased grip aperture (inc) or maximum grip aperture (mga) during the second part of the movement.

#### d. Two additional checks for keypress integrity based on group means

It is again notable that individual correlations between keypress and kinematic MTs were highly variable (Figure S13). Although this might suggest that keypress times were inaccurate indices of movement start and stop times in some participants, on reflection we believe a more likely explanation is that this simply reflects the lack of variability in movement times within blocks, resulting in it being difficult to capture the effects using correlations on an individual level.

As before, a better approach would be to measure the correlation between the mean MTs for all participants across all trials with the comparable mean kinematic MTs. This analysis shows a very high correlation, $r^2 = 0.73$ in the foveal condition and $r^2 = 0.93$ in the peripheral condition, showing a very strong overall correspondence between the two measures (see Figure S14).

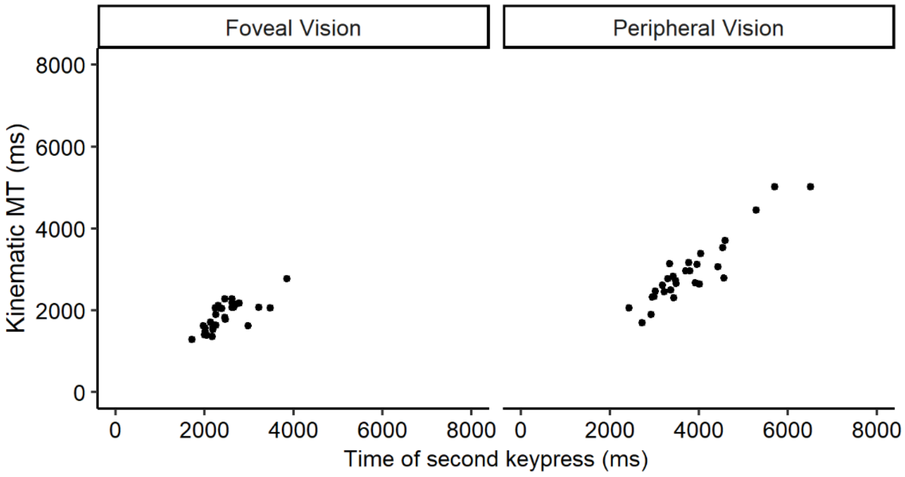


**Figure S14. Correlations between kinematic and keypress MT for each participant across all trials, in each condition.**

A direct comparison of means across the different measures confirms this view. Although from Figure S15, it seems that participants in the peripheral condition pressed slightly later, on average, than would be expected based on the kinematic MT, this error was minimal, and did not alter the overall pattern of results.

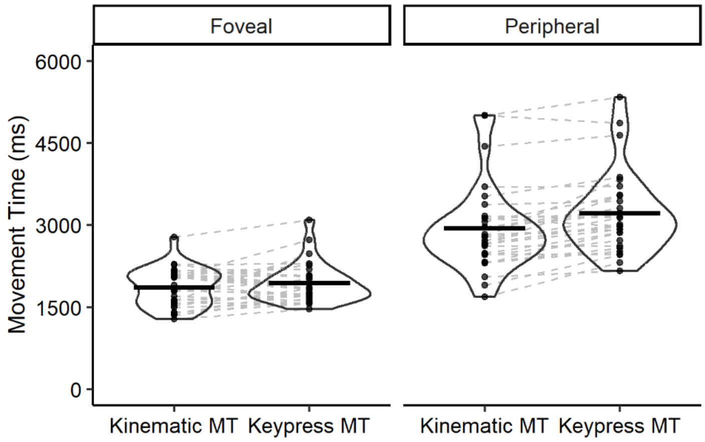


**Figure S15. Comparison between kinematic and keypress movement times in the physical action group.** Conventions as in Figure S4.

### 4. Experiment 3

#### a. Additional analysis on MT

As an additional check of our results, we performed the same extra analyses as for Experiments 1 and 2.

Similar to the analysis in the main manuscript, the best-fitting model included only the main effects of *Action* and *Mime* (Figure S16). A stastistical model including an effect of *Action* fit the data much better than a null model ($\lambda_{adj}$ = (589), with longer movement times in the motor imagery group than in the physical action group. Adding a main effect of *Mime* also improved the fit ($\lambda_{adj}$ = 17), with longer movement times in the mime condition than in the control condition. Contrary to our prediction, adding the interaction *Action* × *Mime* did not improve the fit ($\lambda_{adj}$ = 1.7).

This analysis shows that the participants pressed the key accurately in the physical action group, supporting our experimental design (compare the left panel of Figure 5 from the main manuscript to Figure S16).

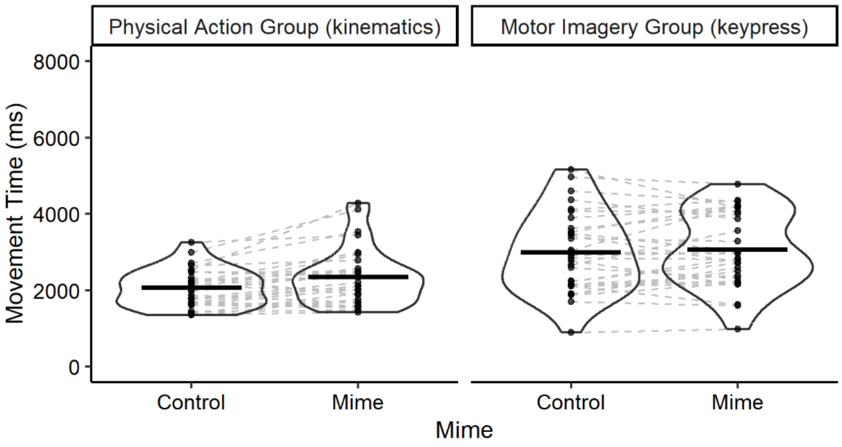


**Figure S16. Effect of varying the quality of online control (Control vs. Mime) on movement times in the physical action and motor imagery groups.** Conventions as in Figure S1.

#### b. Keypress Reaction Times

Neither the Motor-Cognitive model nor the Functional Equivalence view make any specific predictions regarding the RT. We thus conducted an exploratory analysis on the data, presented in Figure S17. The best-fitting model included the main effects of *Action* and *Mime*: participants took longer to initiate an imagined movement compared to the physical action and took longer in the mime condition than the control one. A statistical model including an effect of *Action* fit the data much better than a null model ($\lambda_{adj}$ > 1000), and so did a model including a main effect of *Mime* ($\lambda_{adj}$ = 350). Adding the interaction *Action* × *Mime* did not improve the fit ($\lambda_{adj}$ = .39).

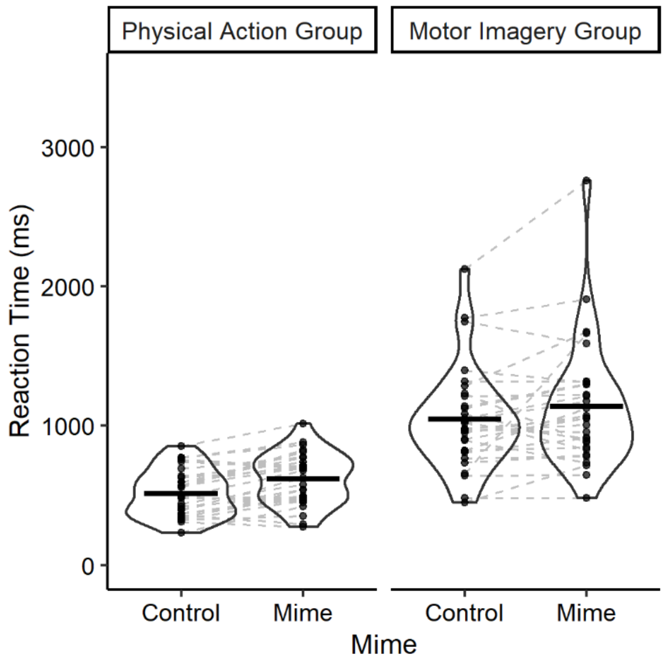


**Figure S17. Effect of varying the quality of online control (Control vs. Mime) on reaction times in the physical action and motor imagery groups.** Conventions as in Figure S2.

#### c. First check for keypress integrity: Individual kinematic correlations

*Methods.* We again measured the correspondence between button-pressing and actual reaction times and movement times measured using the Polhemus motion tracking system, using the same criteria as in Experiments 1 and 2. We applied Pearson correlation for each participant and each *Mime* condition.

*Results.* Moderate correlations were observed between the keypressing indicating the start of the movement and the start of the movement based on the kinematic measure (RT), as well as between the keypressing indicating the end of the movement and the time of the end of the movement given by our three different kinematics markers (MTvel, MTinc and MTmga; Figure S18). Correlations with the velocity-based end of movement were again the highest.

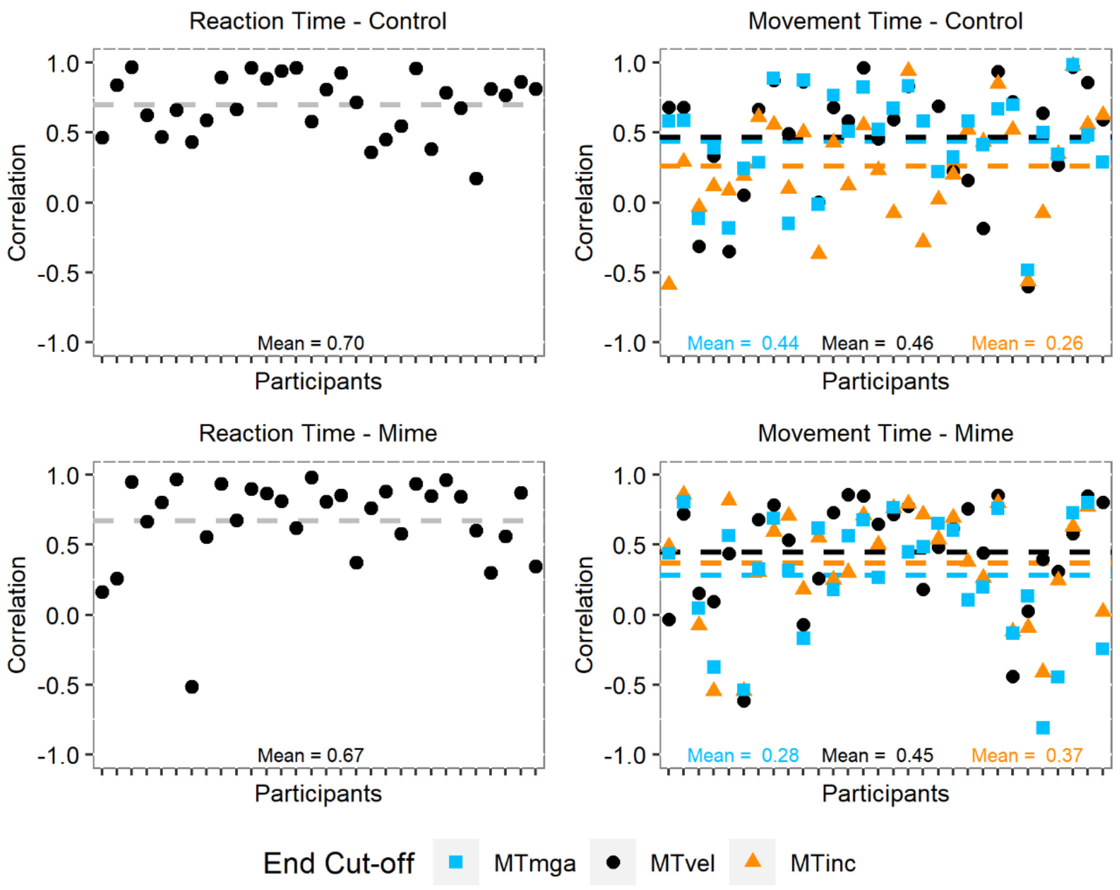


**Figure S18. Pearson correlation values between the keypress data and the kinematics parameters in Experiment 2.** Symbols are individual correlation values for the reaction times (left panel) or the movement times (right panel). In the right panel, MT values on the same column belong to the same participant. Dashed lines indicate the mean correlation for each marker. MT = Movement Time, defined by 3 different criteria: lowest velocity (vel), increased grip aperture (inc) or maximum grip aperture (mga) during the second part of the movement.

#### d. Two additional checks for keypress integrity based on group means

It is again notable that individual correlations between keypress and kinematic MTs were highly variable (Figure S18). Although this might suggest that keypress times were inaccurate indices of movement start and stop times in some participants, on reflection we believe a more likely explanation is that this simply reflects the lack of variability in movement times within blocks, resulting in it being difficult to capture the effects using correlations on an individual level.

As before, a better approach would be to measure the correlation between the mean MTs for all participants across all trials with the comparable mean kinematic MTs. This analysis shows a very high correlation, $r^2 = 0.92$ for the Control and $r^2 = 0.97$ for the Mime, showing a very strong overall correspondence between the two measures (see Figure S19).

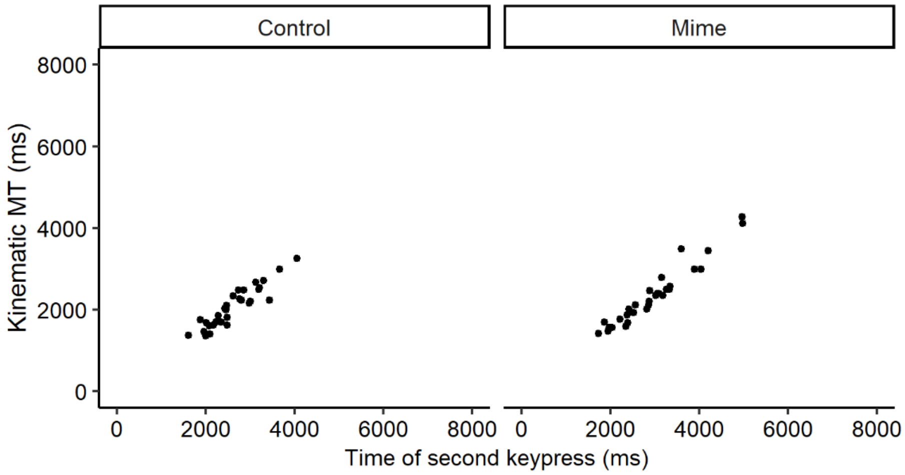


**Figure S19. Correlations between kinematic and keypress MT for each participant across all trials, in each condition.**

A direct comparison of means across the different measures confirms this view. It is clear from Figure S20 that any error in keypress MTs was minimal and not systematic, and did not alter the overall pattern of results.

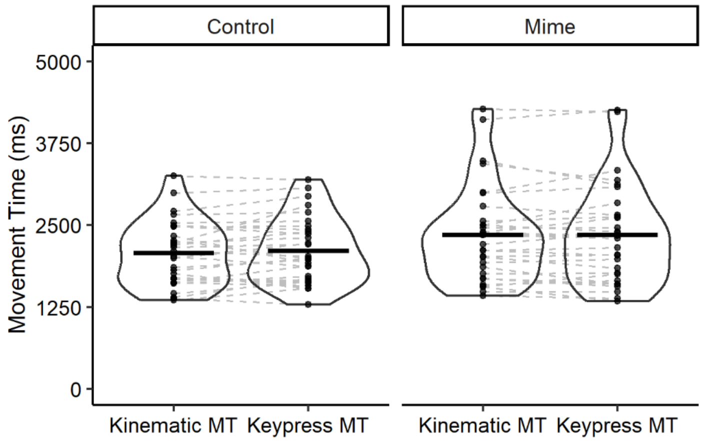


**Figure S20. Comparison between kinematic and keypress movement times in the physical action group.** Conventions as in Figure S4.
